# Engineering Human Pluripotent Stem Cell Lines to Evade Xenogeneic Transplantation Barriers

**DOI:** 10.1101/2023.06.27.546594

**Authors:** Hannah A. Pizzato, Paula Alonso-Guallart, James Woods, Bjarki Johannesson, Jon P. Connelly, Todd A. Fehniger, John P. Atkinson, Shondra M. Pruett-Miller, Frederick J. Monsma, Deepta Bhattacharya

## Abstract

Allogeneic human pluripotent stem cell (hPSC)-derived cells and tissues for therapeutic transplantation must necessarily overcome immunological rejection by the recipient. To define these barriers and to create cells capable of evading rejection for preclinical testing in immunocompetent mouse models, we genetically ablated *β2m*, *Tap1*, *Ciita*, *Cd74*, *Mica*, and *Micb* to limit expression of HLA-I, HLA-II, and natural killer cell activating ligands in hPSCs. Though these and even unedited hPSCs readily formed teratomas in cord blood-humanized immunodeficient mice, grafts were rapidly rejected by immunocompetent wild-type mice. Transplantation of these cells that also expressed covalent single chain trimers of Qa1 and H2-K^b^ to inhibit natural killer cells and CD55, Crry, and CD59 to inhibit complement deposition led to persistent teratomas in wild-type mice. Expression of additional inhibitory factors such as CD24, CD47, and/or PD-L1 had no discernible impact on teratoma growth or persistence. Transplantation of HLA-deficient hPSCs into mice genetically deficient in complement and depleted of natural killer cells also led to persistent teratomas. Thus, T cell, NK cell, and complement evasion are necessary to prevent immunological rejection of hPSCs and their progeny. These cells and versions expressing human orthologs of immune evasion factors can be used to refine tissue- and cell type-specific immune barriers, and to conduct preclinical testing in immunocompetent mouse models.

## Introduction

Pluripotent stem cell (PSC)-based therapies for regenerative medicine are beginning to fulfill their promise, with many new developments in recent years. Some recent examples of PSC-based therapies in clinical trials include neural precursors to treat Parkinson’s disease^1–4^, retinal pigment epithelium for macular degeneration^5–14^, cardiomyocytes for heart failure^15–17^, pancreatic endoderm or islet cells for type 1 diabetes^18,19^, and several cancer immunotherapies^20–23^. Yet immunological rejection of PSC-based therapies remains a limitation to their widespread adoption^24^. To circumvent rejection, these therapies can be combined with systemic immunosuppression, but this leaves patients susceptible to infections and even cancer. Alternatively, induced PSCs (iPSCs) can be used autologously, but this has the practical limitation of scalability. A third option is genetic modification of the starting PSCs to evade immunological rejection. Indeed, this is an area of active research, with many recent studies reporting PSC modifications that evade recognition by specific immune cell types^25–34^. Yet the immunological barriers to transplantation vary substantially between cell types and tissues, and experimental conclusions on the necessary modifications can differ depending on the assays used to test immune evasion. A resource of well-characterized PSCs edited to evade specific components of the immune system would thus be valuable to the community to define cell-type and organ-specific barriers to transplantation and the minimum combinations of edits required to evade these barriers.

T cell-mediated rejection is the best understood immunological barrier to allografts due to mismatched human leukocyte antigen (HLA). HLA is the human specific major histocompatibility complex (MHC) and consists of highly polymorphic cell surface molecules that present peptides to T cells. Class I molecules are expressed on nearly all nucleated cells, while class II is restricted almost exclusively to antigen presenting cells (APCs). These molecules present cytoplasmic (class I) or vesicular (class II) peptides to cytotoxic CD8^+^ and helper CD4^+^ T cells, respectively, and serve as a signal to alert the immune system to pathogens and malignancies. Yet these same beneficial pathways can lead to rejection of allografts, as a very high frequency of the T cell repertoire recognizes antigens from HLA-mismatched donors^35^.

For the purposes of PSC-based allogeneic transplantation, genetic ablation of molecules required for HLA expression might render cells invisible to direct recognition by T cells. For example, β2-microglobulin (β2m) is the common light chain of MHC-I, and its mutation leads to MHC-I deficiency^36,37^. Similarly, ablation of transporter associated with antigen processing 1 (TAP1), a subunit of the transporter responsible for delivering peptides into the endoplasmic reticulum (ER) for MHC-I loading, sharply reduces MHC-I surface expression^38^. Mutation of CD74 (invariant chain, Ii), a polypeptide that shuttles MHC-II to the endosome to bind its antigenic peptides, causes ∼100-fold decrease in MHC-II cell surface expression^39^. Similarly, deficiency in the transcription factor class II transactivator (CIITA) leads to a nearly complete absence of MHC-II expression in most cell types^40,41^.

While genetic removal of HLA would likely be necessary for the generation of ‘universal’ donor PSCs, it is unlikely to be sufficient. Mice reject skin allografts deficient of MHC almost as rapidly as transplants with intact mismatched MHC^42,43^. This demonstrates that graft rejection is not solely mediated by direct MHC:T cell interactions. As one mechanism, MHC-I normally engages inhibitory receptors on natural killer (NK) cells^44–47^. If activating receptors are then also engaged by their cognate ligands, NK cells can kill their target cells. Yet specific knowledge of these pathways offers opportunities to intervene and edit PSCs to substantially reduce NK cell recognition. The activating receptor NK group 2 member D (NKG2D) is expressed on nearly all NK cells, and its inhibition reduces cytotoxicity by ∼50%^48,49^. Thus, genetically ablating ligands for NKG2D, such as MHC class I chain-related proteins A and B (MICA and MICB)^50,51^, is expected to substantially dampen NK cell-mediated cytotoxicity. Reciprocally, HLA-E and HLA-G are ligands for the human NK cell inhibitory receptors NK group 2 member A (NKG2A) and immunoglobulin-like transcript 2 (ILT2)/killer cell immunoglobulin-like receptor, two immunoglobulin domains and long cytoplasmic tail 4 (KIR2DL4), respectively^52–59^. These nonclassical HLA molecules are not polymorphic and present a limited pool of self peptides. Expression of these molecules is also expected to attenuate NK cell activation and cytotoxicity. Still, NK cells alone cannot explain the rapid rejection of mouse skin allografts^42^, suggesting the contribution of still other immune pathways.

Graft-reactive antibodies, which often bind foreign HLA molecules, and subsequent deposition of complement also contribute to rejection^60–63^. Complement deposition directly lyses target cells and activates phagocytes, B cells, and other lineages of the immune system. Though the antibody-mediated classical complement pathway receives the most attention in organ transplants, the antibody-independent alternative and lectin pathways likely also contribute to graft injury and rejection^64–68^. Because the complement system can damage normal tissues, there are several endogenous inhibitors of complement deposition, for which deficiencies cause diseases such as subsets of hemolytic anemia^69^ and age-related macular degeneration^70^. These include CD46, or membrane cofactor protein (MCP), which promotes cleavage of cell bound C3b and C4b^71–73^; CD55, or decay-accelerating factor (DAF), which disrupts the formation of the C3 and C5 convertases^74^; and CD59, which prevents the assembly of the membrane attack complex and subsequent cell lysis^75,76^. Complement receptor 1 (CR1) is an additional inhibitor of complement activation, which may inhibit complement deposition more rapidly and efficiently than CD46 and CD55^77^, but has a much more limited cellular distribution^78^. Overexpression of these molecules on PSCs would thus be expected to attenuate complement-mediated rejection.

Finally, phagocytes have also been shown to play a role in graft rejection^79–83^. Macrophages phagocytose damaged and dead cells as well as foreign cells^84^. In the setting of transplant rejection, inflammation triggers phagocytic activity as a mechanism to clear foreign pathogens and dying cells^85^. Not only can phagocytosis remove donor cells, it can also serve to indirectly prime T cells to donor peptides. As with NK cells and the complement system, endogenously expressed factors limit unchecked phagocytosis. For example, CD47 is an immunoglobulin-like protein that interacts with signal regulatory protein α (SIRPα) on phagocytes to inhibit phagocytosis^86,87^. In this sense, CD47 acts as a “don’t eat me” signal and can promote survival of allografts and tumors^27,29,31–33,88^. Similarly, programmed death-ligand 1 (PD-L1) directly inhibits T cell activation by binding to programmed cell death 1 (PD-1) but has also been found to inhibit antigen presentation and phagocytosis by macrophages^89^. Recent studies have demonstrated that overexpression of PD-L1 alone or in combination with CD47 increases the survival of PSC or PSC-derived transplants^27,34^. Finally, CD24 can also inhibit phagocytosis by binding to the inhibitory receptor sialic-acid-binding Ig-like lectin 10 (Siglec-10) on macrophages^90^. Enforced expression of these molecules is thus expected to improve allogeneic PSC-graft survival.

To create resources to define immunological barriers for specific cell types or organs, we have generated a series of genetic modifications of the well-characterized H1 human embryonic stem cells (hESCs)^91^, engineered to evade immune recognition by each of the pathways discussed above. H1 cells devoid of HLA-I and -II and the NK cell activating ligands MICA and B, and additionally expressing inhibitory proteins for NK cells, complement, and phagocytes can cross xenogeneic barriers and form teratomas in wild-type mice. By testing combinations of these immune evasion factors, we have also demonstrated that inhibition of both complement and NK cells are required for graft persistence, while inhibition of phagocytosis is dispensable. These results present a strategy for engineering ‘universal’ hPSCs that could be used for clinically relevant transplants.

## Results

### Generation of an HLA-I/II and MICA/B-deficient hESC line

We designed a CRISPR/Cas9 workflow to ablate 6 genes involved in T cell and NK cell recognition: *β2m* and *Tap1* to eliminate HLA-I and evade CD8^+^ T cells^92–94^; *Cd74* and *Ciita* to prevent HLA-II expression and CD4^+^ T cell recognition^95–97^, and *Mica* and *Micb* to evade activating NKG2D receptors on NK cells^48–51^. Although individual mutations in *β2m* and *Ciita* likely eliminate all relevant HLA-I/II expression, prior studies in mice have suggested that residual MHC expression can still be observed in some knockout cell types^94,95,97–102^. In these cases, further ablation of *Tap1* and *CD74* would likely ensure the absence of physiologically relevant surface HLA-I/II expression. H1 (wild-type) hESCs were iteratively nucleofected with a Cas9 construct and up to 3 gRNA-encoding vectors to target each gene. The pool of transfectants was then subjected to next generation MiSeq analysis to quantify the frequencies of frameshift mutations in each targeted gene. Afterwards, individual colonies were picked and sequenced to identify those that carried frameshift mutations. Individual cells from these colonies were then single cell-sorted to generate clones. Another round of sequencing was performed to verify the mutations and confirm the absence of mosaicism. Clones were then expanded and karyotyped. The overall workflow is depicted in **Figure 1A**. Through this workflow, we generated a karyotypically normal hESC line that carries frameshift mutations in all alleles of *β2m*, *Tap1*, *Cd74*, *Ciita*, *Mica*, and *Micb* (**Table S1**). For *Mica* and *Micb*, an additional round of targeting was required to ablate an in-frame and potentially functional fusion protein (**Figure S1B**). Through this process, we generated lines that are HLA-I/II-deficient (HLA-I/II-KO) and HLA-I/II and MICA/B deficient (HM-KO), summarized in **Table S2**.

**Figure 1.**
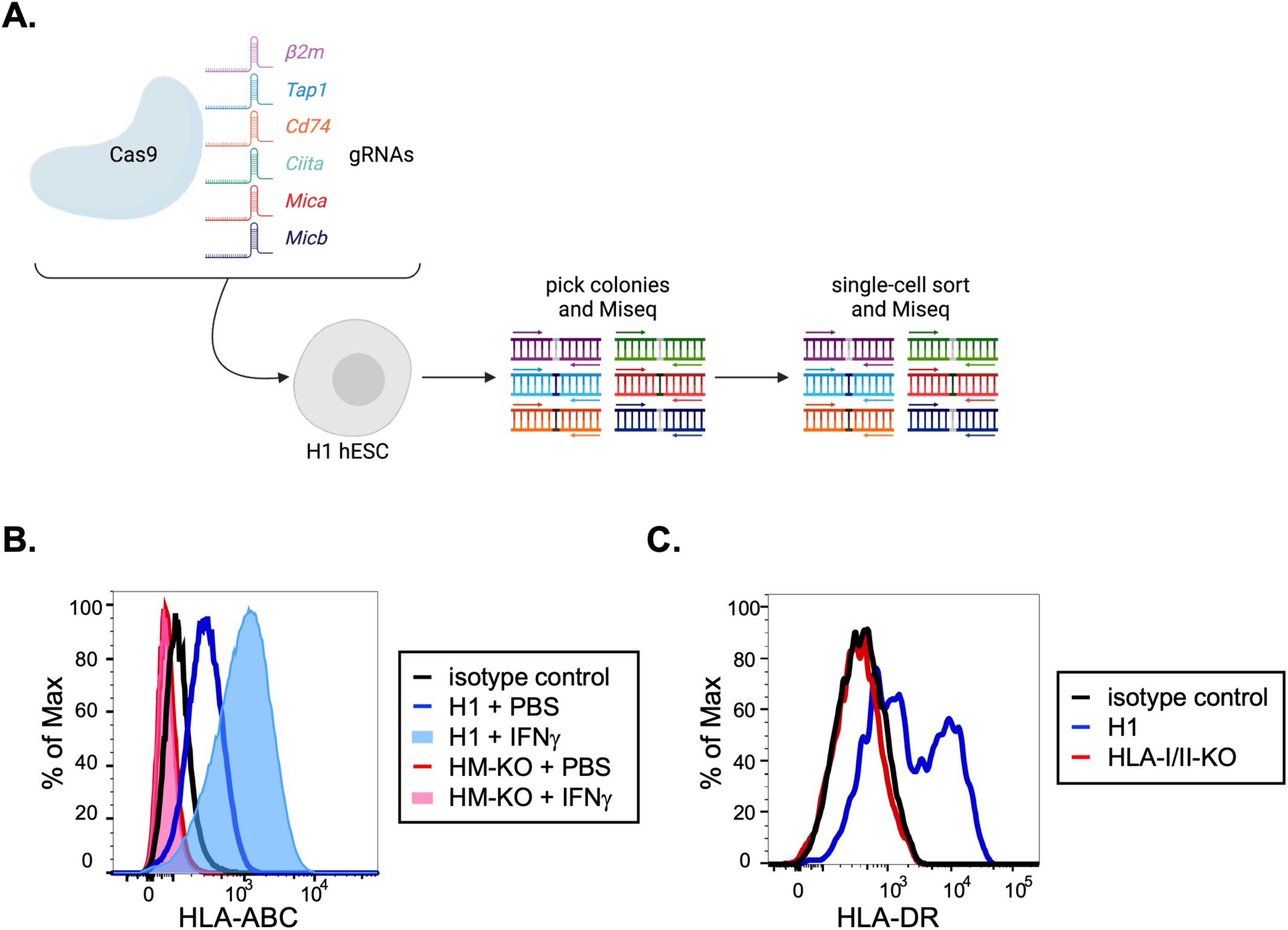
Generation of an HLA-I/II and MICA/B-deficient hESC line. (A) Schematic representation of Cas9 editing strategy. Wild-type H1 hESCs were transfected with a Cas9 construct and up to 3 gRNAs at a time. Guide RNAs for *β2m*, *Tap1*, *Cd74*, *Ciita*, *Mica*, and *Micb* were used. Following transfection, individual hESC colonies were picked and sequenced to identify colonies with frameshift mutations in the target genes. Colonies with frameshift mutations were then expanded, clonally sorted, and re-sequenced for frameshift mutations. The resulting cells were then iteratively targeted by gRNAs to generate a cell line in which all alleles carried frameshift mutations. (B) Representative flow cytometric histograms of HLA-ABC (HLA-I) expression by HM-KO and parental H1 hESCs treated with PBS as a vehicle control or 20ng/mL IFNγ for 24hrs. Additionally, cells were stained with an isotype control antibody to account for background staining. (C) Representative histograms of HLA-DR (HLA-II) expression of dendritic cell (DC)-like cultures differentiated from H1 or HLA-I/II-KO hESCs. hESCs were differentiated into hematopoietic progenitors using the STEMdiff Hematopoietic Kit (STEMCELL Technologies) and then were further differentiated into DC-like cells with 100ng/mL GM-CSF and 100ng/mL IL-4.

Undifferentiated hESCs express very low levels of HLA-I and do not express HLA-II^103^. Therefore, to confirm HLA-I deficiency, wild-type H1 hESCs and HM-KO hESCs were treated for 24hrs with 20ng/mL interferon (IFN) γ to induce HLA-I expression^104,105^. The HM-KO cells lacked HLA-I expression, whereas their wild type counterparts significantly upregulated expression upon IFNγ treatment (**Figure 1B**). To confirm ablation of HLA-II, hESCs were sequentially differentiated into hematopoietic progenitors, dendritic cell (DC)-like cells via treatment with 100ng/mL granulocyte-macrophage colony-stimulating factor (GM-CSF), and activated DCs through the addition of 100ng/ml IL-4^106^. These experiments confirmed that HLA-II was not detectably expressed by HLA-I/II-KO hESC-derived DC-like cells (**Figure 1C**).

During the course of this work, several reports raised concerns that the use of CRISPR/Cas9 tended to favor cells with mutations in the proto-oncogene *Tp53*^107–109^. In light of these findings, we performed whole exome sequencing on our KO lines. After the first round of edits to generate the HLA-I-KO line, we detected a CGG to CAG mutation in *Tp53*, creating a missense R248Q change (**Figure S1B**). This dominant negative R248Q change is one of the most common mutations in human cancers^110^. No other potentially oncogenic or pathogenic sequence variants were observed, as defined by the National Center for Biotechnology Information’s ClinVar database^111^. Fortuitously, the point mutation in *Tp53* created a *de novo* PAM sequence and gRNA site, allowing for selective reversion of the sequence. Single-stranded donor oligonucleotides were designed such that the mutant CAG codon was reverted to AGG, which restores arginine but can be distinguished from the other wild-type allele by next generation sequencing (**Figure S1B**). These donor oligonucleotides were transfected into HM-KO hESCs alongside a gRNA-Cas9 ribonucleic acid-protein complex. We screened by MiSeq and selected two clones that contained both AGG and CGG codons, thereby functionally restoring the wild-type p53 amino acid sequence.

### Humanized mice are incapable of rejecting allogeneic teratomas

*In vitro* studies generally fail to fully capture the complexity of mechanisms that lead to transplantation rejection. Therefore, to test human immune evasion by these cells *in vivo*, we generated humanized mice. NOD.Cg-*Kit^W-41J^ Tyr* ^+^ *Prkdc^scid^ Il2rg^tm1Wjl^*/ThomJ (NSG-W41) mice lack B, T, and NK cells and carry a mutation in *Kit* that allows robust, long-term engraftment of human HSCs without irradiation^112–115^. We transplanted 1 − 10^5^ human cord blood CD34^+^ cells into unconditioned NSG-W41 mice. At least two months post-transplantation, we typically observed ∼3% human chimerism in the blood and ∼50% in the spleen, with robust B and T cell chimerism (**Figures 2A and B**).

**Figure 2.**
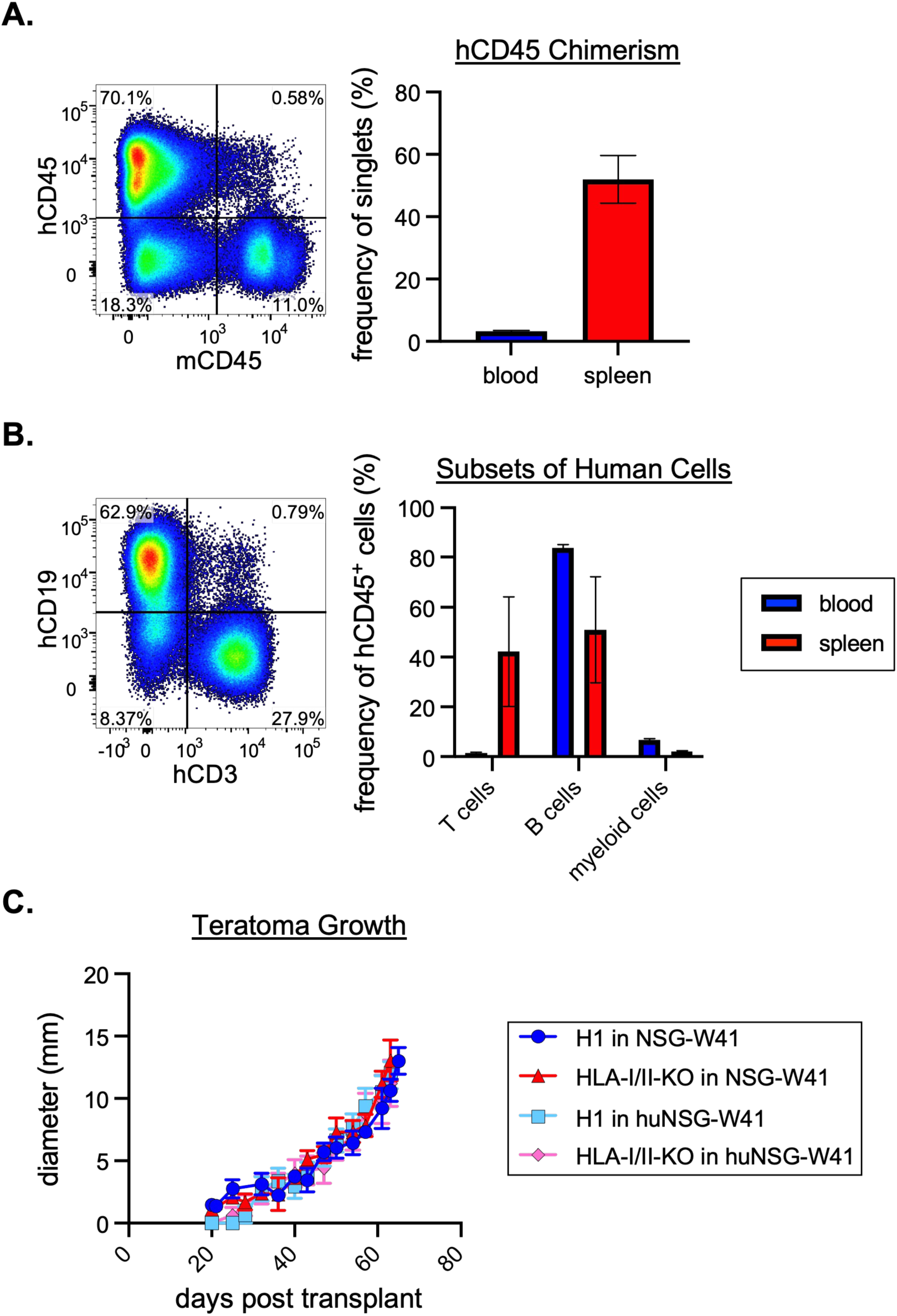
Humanized mice are incapable of rejecting allogeneic teratomas. (A) Donor chimerism of cord blood-humanized NSG-W41 (huNSG-W41) mice. 1 − 10^5^ human cord blood CD34^+^ cells were transplanted into unconditioned NSG-W41 mice. At 2+ months post-transplantation, mice were bled to confirm human chimerism via hCD45 staining. Splenic chimerism was determined for a subset of mice. A representative flow cytometry plot of splenic human chimerism gated on live single cells (left). Blood and splenic chimerism are displayed as the mean values ± SEM of 63 and 5 mice, respectively (right). (B) Cellular composition of human cells in huNSG-W41 mice. The human population of cells from blood and splenic samples were analyzed for the frequency of T cells (CD3), B cells (CD19), and myeloid cells (CD13). A representative flow cytometry plot of splenic T and B cells gated on live single cells is shown in the left panel. Mean values ± SEM are shown for the blood (63 mice) and spleen (5 mice) (right). (C) Teratoma assay of H1 and HLA-I/II-KO hESCs in NSG-W41 and huNSG-W41 mice. 1 − 10^6^ hESCs were embedded in Matrigel and injected subcutaneously into the hind flank of a mouse. Mean values ± SEM are shown for each group of mice. Five to ten mice are represented per group from two independent experiments. P values greater than 0.05 by 2-way ANOVA with post-hoc Tukey’s multiple comparisons test are not depicted.

Using these humanized mice, we performed *in vivo* teratoma assays to test immune evasion by our cells^116^. Persistence and expansion of teratomas can be taken as evidence of immune evasion *in vivo*. One million wild-type H1 or HLA-I/II-KO hESCs were embedded in Matrigel and injected subcutaneously in NSG-W41 or humanized NSG-W41 (huNSG-W41) mice. Unexpectedly, unmodified H1 cells were readily able to form teratomas in humanized mice at a similar rate and size to those in unhumanized NSG-W41 mice (**Figure 2C**). These data suggest that cord blood-humanized mice are not reliable representatives of human immune-mediated transplant rejection. Indeed, some aspects of the immune response in humanized NSG-W41 mice are not fully functional, such as NK cells^117^ and antibody responses^118^. Though it is possible that other humanized systems might have performed better, each of these models has deficiencies ranging from MHC/HLA mismatches between thymic epithelial cells that educate T cells and peripheral antigen presenting cells, lack of binding by mouse cytokines to human receptors, and graft versus host disease when mature T cells are transferred^117–120^. Given the failure of humanized mouse models to reject even wild-type human teratomas, we sought to develop more stringent systems to test rejection in immunocompetent wild-type C57BL/6 (B6) recipients. These animals have fully intact xenogeneic immune barriers, and, as such, represent an exceptionally high bar for transplantation^121–132^. Success in this system would raise confidence that cells could also cross allogeneic barriers.

### Immune evasion construct design and expression

MHC-deficient grafts are still rapidly rejected when transplanted to allogeneic immunocompetent mouse recipients^42,133^. Thus, we considered it unlikely that HLA-I/II-KO hESCs would evade rejection in wild-type xenogeneic mouse recipients without further modifications. Other studies have revealed additional mediators of rejection to be natural killer (NK) cells^134^, complement^135^, and phagocytes^88^. However, the depletion or inhibition of these individually is not enough to prevent graft rejection, although it does prolong graft survival^136–138^. To inhibit all these pathways, we designed a series of constructs that encode mouse and human orthologs of immune evasion factors including CD46, CD55, and CD59 for complement; HLA-E and HLA-G for NK cells; and CD47 for phagocytosis (summarized in **Table S3**). Mice have orthologs of CD47, CD55 and CD59, but Crry in mice is the functional homologue of human CD46^139–142^. The mouse ortholog for HLA-E is the nonclassical MHC molecule Qa1^143,144^. There is no mouse ortholog for HLA-G, but H2-K^b^ inhibits Ly49C^+^ NK cells^145^, which are unique to mice. MHC and HLA molecules can be covalently linked to a peptide and β2m to form single chain trimers^56,59,146,147^. These trimers cannot exchange their peptide or β2m with endogenous HLA^148^, and thus cannot rescue HLA deficiencies.

We designed two different types of constructs to test the efficacy of these immune evasion genes. For rapid testing of the mouse immune evasion genes, we generated lentiviral constructs (an example construct is shown in **Figure 3A**). This system allows for testing of HM-KO derivatives that express different combinations of factors. Each lentivirus construct encodes for a single mouse immune evasion factor, with expression driven by the ubiquitin (UBC) promoter^149,150^. The constructs link the immune evasion gene by a T2A sequence to green fluorescent protein (GFP), which, alongside flow cytometric confirmation of surface expression, allows for facile identification of transduced cells.

**Figure 3.**
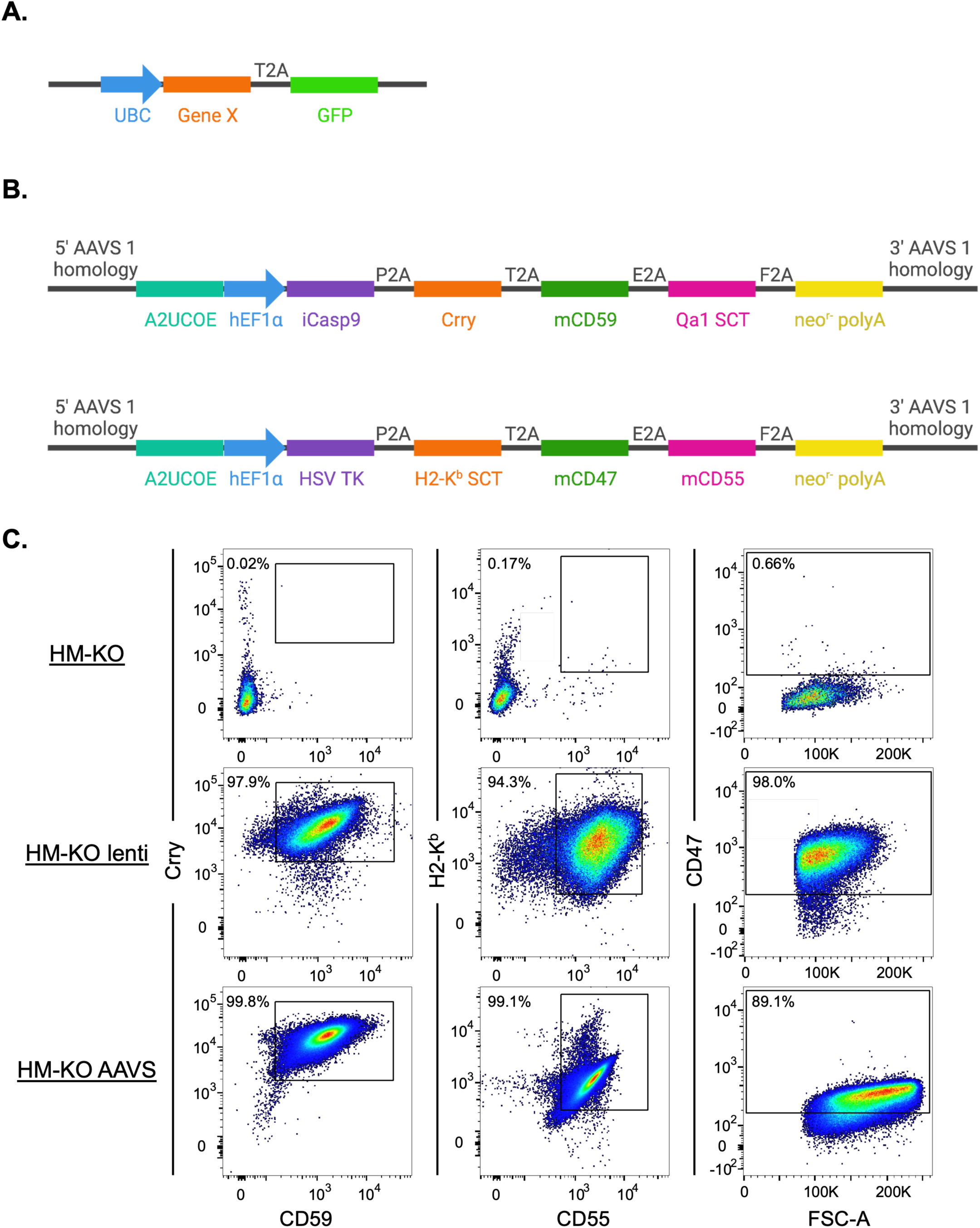
Mouse immune evasion construct design and expression. (A) An example lentivirus construct for expression of mouse immune evasion genes. Expression is driven by the ubiquitin (UBC) promoter. Green fluorescent protein (GFP) is also expressed and linked to the upstream immune evasion Gene X via a T2A sequence. (B) Two AAVS constructs encode a panel of mouse immune evasion genes. Each construct contains an A2UCOE insulator element, and expression is driven by the human elongation factor 1α (hEF1α) promoter. The immune evasion genes Crry, mCD59, Qa1 single chain trimer (SCT), H2-K^b^ SCT, mCD47, and mCD55 are linked by 2A sequences. A suicide gene [inducible caspase 9 (iCasp9) or Herpes Simplex virus (HSV) thymidine kinase (TK)] is present in each construct, as well as a neomycin resistance gene. The constructs have 5’ and 3’ AAVS1 targeting homology arms. (C) Flow cytometric analysis of mouse inhibitory protein expression on HM-KO cells, as well as HM-KO cells transduced with all the lentiviral constructs (HM-KO lenti) or transfected with both mouse AAVS constructs (HM-KO AAVS).

We also designed adeno-associated virus site 1 (AAVS1) constructs encoding either mouse or human immune evasion proteins (**Figures 3B and S2A**, respectively) that target a specific integration site in the PPP1R12C locus. This locus is often referred to as a “safe harbor” for heterologous gene expression into the human genome^151^. Each construct also contains an inducible suicide gene [Herpes Simplex virus (HSV) thymidine kinase (TK) or inducible caspase 9 (iCasp9)], which could be used to eliminate engrafted cells if needed (with treatment with ganciclovir or AP1903, respectively^152–154^), and a drug resistance gene to allow for selection of stable transfectants. Expression of the immune evasion factors is driven by the human elongation factor 1α (hEF1α) promoter^155^, and genes are linked by 2A sequences^156–162^.

We transduced HM-KO hESCS with either all the lentiviral constructs encoding mouse orthologs of immune evasion factors (HM-KO lenti) or transfected them with the AAVS constructs encoding either the mouse (HM-KO AAVS) or human immune evasion factors. We confirmed expression of the proteins and purified cells expressing all the proteins using fluorescence activated cell sorting (FACS) (**Figures 3C and S2B**). Of note, HM-KO lenti cells had slightly higher expression of proteins than did HM-KO AAVS cells.

The lentiviral transduction system was successful with only modest silencing observed over time with passages. Yet for our AAVS-targeting constructs, after multiple rounds of drug selection and sorting, both in bulk and as clones, we repeatedly observed silencing of our constructs, even when cells were maintained under neomycin selection (**Figure S3A**). Most AAVS1 constructs target a region downstream of the actual AAV integration site, which can become highly methylated and is thus prone to silencing^163^. Therefore, we generated new immune evasion constructs and gRNAs to target the construct to the correct AAV integration site. We also included an *HNRPA2B1-CBX3* ubiquitously acting chromatin opening elements (A2UCOE) insulator to further help prevent silencing^164^. These constructs allowed for the stable expression of the immune evasion factors over time (**Figure S3B**).

### HM-KO lenti and AAVS hESCs evade immune rejection in wild-type mice

To quantify the ability of these edited cells to evade rejection, we performed teratoma assays in B6 mice using parental H1, HM-KO, HM-KO lenti, and HM-KO AAVS hESCs. While detectable HM-KO teratomas persisted slightly longer than did H1 teratomas, both were ultimately rejected (H1s within 3wks, and HM-KOs within 7wks; **Figure 4A**). Remarkably, both HM-KO lenti and AAVS teratomas persisted for at least 23wks in these immunocompetent, xenogeneic recipients. The HM-KO lenti and AAVS lines grew and persisted similarly *in vivo*, suggesting the gene delivery system does not affect immune evasion. HM-KO lenti and AAVS teratomas were harvested 102 days after transplantation, sectioned, and stained for DAPI, human mitochondria, and the mouse immune evasion factors, demonstrating the teratomas were in fact human donor cells expressing inhibitory proteins (**Figure 4B**). We found this method of confirming donor cell origin more reliable than bioluminescence due to variable rates of luciferase silencing by our cells in culture. Nonetheless, while the engrafted HM-KO lenti and AAVS cells persisted, teratoma formation was stalled at ∼2mm in diameter (**Figure 4A**).

**Figure 4.**
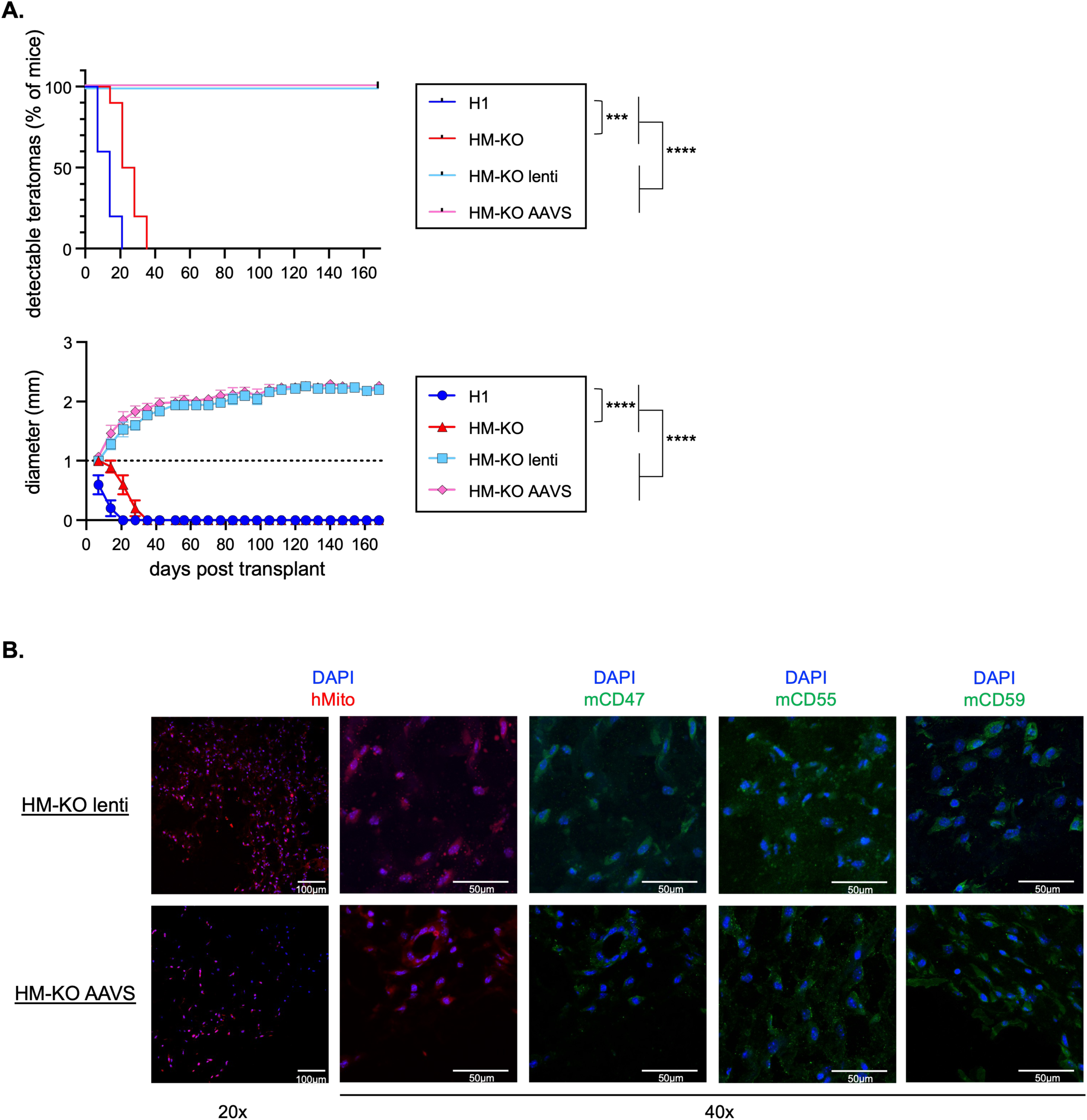
HM-KO lenti and AAVS hESCs evade immune rejection in wild-type mice. (A) Teratoma assay of H1, HM-KO, HM-KO lenti, and HM-KO AAVS cells in B6 mice. Teratomas >1mm in diameter were scored as detectable (top panel), while measurements of the teratoma sizes are depicted in the bottom panel. The dashed line at 1mm represents the limit of detection. Mean values + SEM are shown for each group of mice. Ten mice are represented per group from two independent experiments. For teratoma detection, ***p<0.001 and ****p<0.0001 by log-rank test. For teratoma growth, ****p<0.0001 by 2-way ANOVA with post-hoc Tukey’s multiple comparisons test. (B) Representative immunohistochemistry (IHC) staining of sections of HM-KO lenti and AAVS teratomas removed on day 102 post transplantation into B6 mice. Sections were imaged at 20x magnification with 100µm scale bars and 40x magnification with 50µm scale bars. Sections were stained with DAPI in combination with antibodies for human mitochondria (hMito) or mouse CD47, CD55, and CD59.

Given the plateauing of teratoma growth, we explored whether additional immune barriers remain. We treated mice with CD4, CD8, CD20, or NK1.1 depleting antibodies prior to transplanting hESCs and weekly throughout the duration of the teratoma assay. Efficient cell depletion of CD4^+^ T cells, CD8^+^ T cells, B cells, and NK cells was achieved within 4 days of the first antibody injection (**Figure S4A**). Nonetheless, none of these depletions further promoted the growth of teratomas compared to isotype controls (**Figure S4B**). These data suggest that immune lineage-derived factors may directly or indirectly lead to the differentiation, rather than rejection, of transplanted hESCs *in vivo*, thereby halting teratoma growth.

Given the unreliability of the humanized mouse model system, we validated the human orthologs of these immune evasion genes and constructs *in vitro.* Undifferentiated hESCs express high endogenous levels of complement inhibitors and are themselves resistant to lysis^165–167^. Therefore, to validate the human AAVS constructs for inhibition of complement, we utilized Chinese hamster ovary (CHO) cells, in which such assays are usually performed^168^. CHO cells stably expressing human immune evasion constructs were incubated with anti-CHO antibody, then human C7-deficient serum, and finally were stained for complement fragment deposition. Expression of hCD55 and hCD46 decreased C3c, C3d, and C4c deposition as compared to cells in the same cultures that did not express these proteins (**Figures S5A-C**). The isolated effects of CD59 were more difficult to assess, as it inhibits steps downstream of CD55 and CD46^169,170^.

Undifferentiated hESCs are also resistant to NK cell recognition and lysis^105^. Therefore, we utilized K562 cells, which are highly susceptible to NK cells, to test the efficacy of the inhibitory proteins encoded by our immune evasion constructs. Using flow cytometry, we verified protein expression on K562s (**Figure S5D**) and sorted cells that stably expressed these proteins for downstream assays. To confirm efficacy of the NK cell inhibitory ligands, we incubated unmodified K562s or stably-transfected K562s with human peripheral blood mononuclear cells (PBMCs). While unmodified K562s triggered robust NK cell degranulation, as apparent by CD107a surface expression, HLA-E expressing cells specifically inhibited NKG2A-expressing NK cell degranulation (**Figures S5E and F**). These *in vitro* assays demonstrate the proteins encoded by the human constructs are functional.

### Inhibition of both complement and NK cells is necessary to prevent immunological rejection

Given the extensive modifications to our cells, we performed experiments to define which modifications were necessary and sufficient to evade rejection. Using our mouse lentivirus constructs, we generated lines expressing only the complement, NK cell, or phagocytosis inhibitory factors. We also tested several additional factors to inhibit phagocytosis and an alternative inhibitor for complement. For phagocytosis, we made lentivirus constructs for mouse PD-L1 and CD24 and transduced them into HM-KO cells (**Figure S6**). Due to the large size of the complement inhibitor CR1 and poor packaging in lentiviruses, we generated an AAVS construct to transfect into HM-KO cells (**Figure S6**).

Teratoma assays were performed on all the lines inhibiting individual pathways (**Table S2**) to determine which could reduce immunogenicity and evade rejection similarly to the HM-KO AAVS line. None of these new lines generated and maintained teratomas similarly to the HM-KO AAVS line (**Figure 5A**). However, relative to HM-KO teratomas, HM-KO comp, NK, and CR1 comp teratomas each detectably persisted for longer (p<0.05) and showed increased growth (p<0.001). HM-KO phago and CR1 teratomas did not provide any significant additional evasion relative to HM-KO teratomas. Given the slight advantage that complement and NK cell inhibitory proteins provided to HM-KO cells, we tested whether the combined inhibition of both pathways would achieve HM-KO AAVS levels of evasion. Indeed, HM-KO cells expressing immune evasion factors for both NK cells and complement mirrored the persistence and growth of HM-KO AAVS teratomas (**Figure 5B**). These results suggest that both complement and NK cells are major drivers of rejection in our system, and their inhibition is necessary for the persistence of teratomas. Additionally, both the HM-KO comp and CR1 comp, as well as the HM-KO NK and comp and CR1 NK and comp, had similar persistence and growth, suggesting that CR1 can be a surrogate for Crry and CD55. HM-KO CR1 comp teratomas persisted longer and grew larger than HM-KO CR1 teratomas (p<0.05), suggesting that CD59 provides additional immunological protection.

**Figure 5.**
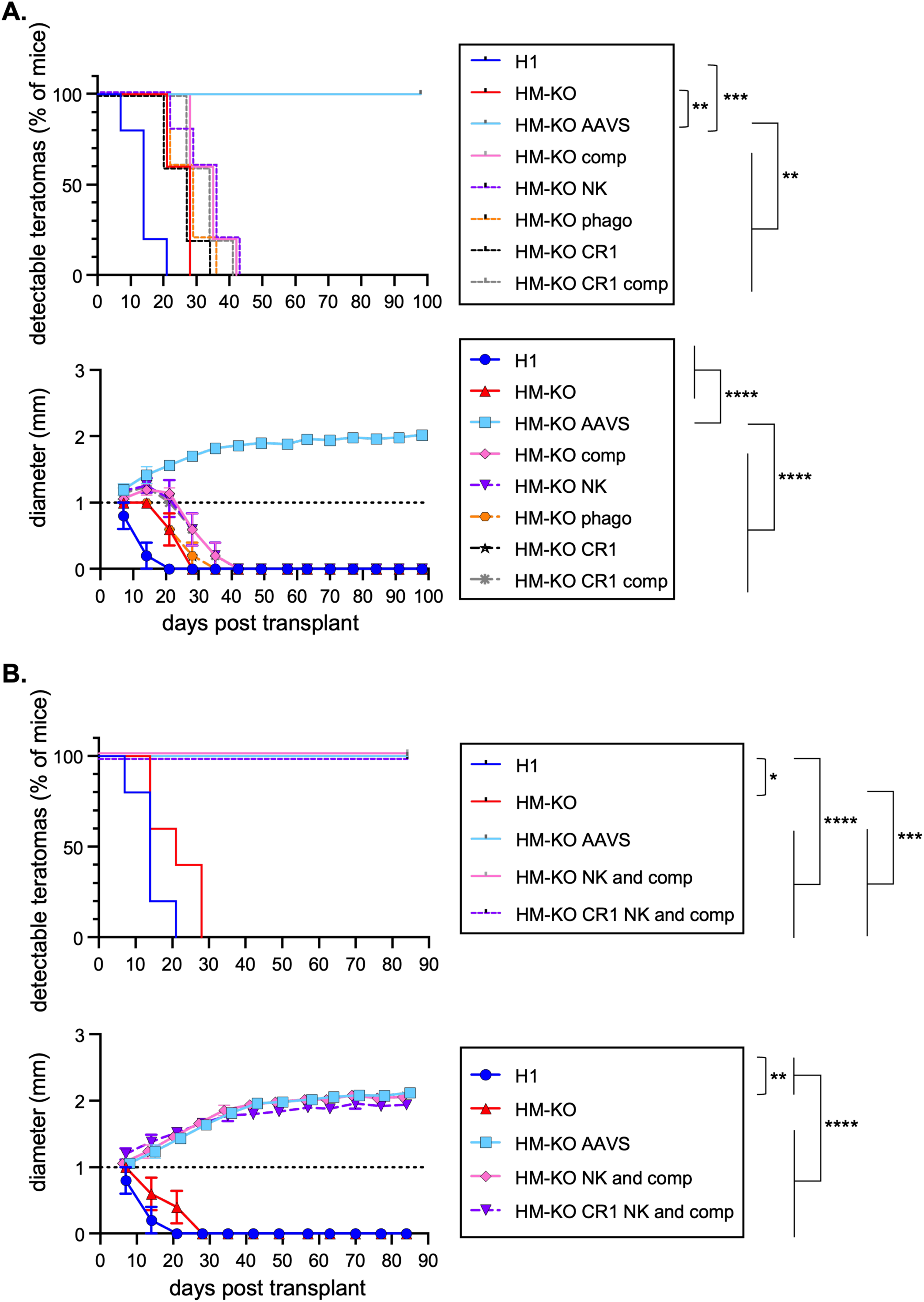
Inhibition of both complement and NK cells is necessary to prevent immunological rejection. (A) Teratoma assay of H1, HM-KO, HM-KO AAVS, as well as HM-KO cells expressing individual groups of immune evasion factors (outlined in Table S2) in B6 mice. The frequency of detectable teratomas is shown in the top panel, while size of the teratomas is depicted in the bottom panel. The dashed line at 1mm represents the limit of detection. Mean values + SEM are shown for each group of mice. Five mice are represented per group. For teratoma detection, **p<0.01 and ***p<0.001 relative to HM-KO AAVS by log-rank test. For teratoma growth, ****p<0.0001 relative to HM-KO AAVS by 2-way ANOVA with post-hoc Tukey’s multiple comparisons test. (B) Teratoma assay of H1, HM-KO, HM-KO AAVS, as well as HM-KO cells expressing both NK and complement inhibitory proteins (outlined in Table S2) in B6 mice. The frequency of detectable teratomas is shown in the top panel, while size of the teratomas is depicted in the bottom panel. The dashed line at 1mm represents the limit of detection. Mean values + SEM are shown for each group of mice. Five mice are represented per group. For teratoma detection, *p<0.05, ***p<0.001, and ***p<0.001 by log-rank test. For teratoma growth, **p<0.01, and ****p<0.0001 by 2-way ANOVA with post-hoc Tukey’s multiple comparisons test.

### HM-KO cells persist in mice deficient in complement and NK cells

The immune evasion genes used above might potentially impact more than just the target pathways. To further investigate the importance of inhibiting both complement and NK cells in orthogonal assays, we measured teratomas using HM-KO cells in complement deficient *C3^−/−^* mice^171^, with or without NK cell depletion. For this experiment, NK cells were depleted using an NK1.1 depleting antibody prior to transplantation of HM-KO cells and weekly throughout the duration of the experiment. Peripheral NK cells were depleted within 4 days of the initial depleting antibody injection (**Figure S4A**). Consistent with HM-KO NK and comp teratomas, both HM-KO teratomas in NK cell-deficient *C3^−/−^* mice and HM-KO NK teratomas in *C3^−/−^* mice persisted for at least 84 days post-transplant (**Figure 6**). While these teratomas still plateaued in diameter, the size was slightly greater than seen with HM-KO AAVS and lenti teratomas in wild-type mice (compare **Figure 6** with **Figure 4A**). As expected, HM-KO teratomas in NK-deficient B6 mice or in isotype control *C3^−/−^* mice mirrored HM-KO NK teratomas and HM-KO comp teratomas in B6 mice (**Figures 5A and 6**). These results confirm that HM-KO cells persist with the inhibition or ablation of both complement and NK cells, while inhibition of phagocytosis is neither necessary nor sufficient.

**Figure 6.**
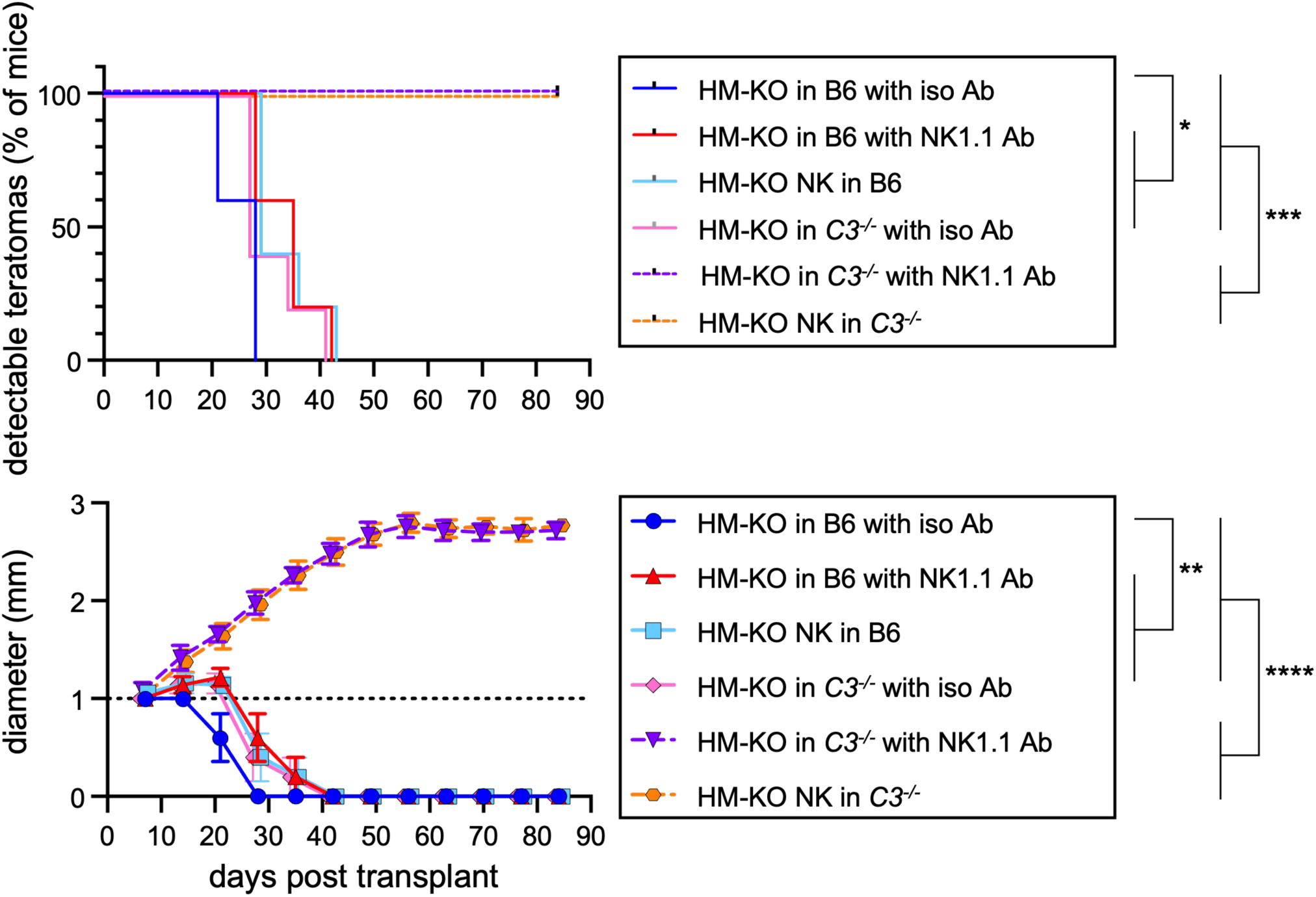
HM-KO cells persist in mice deficient in complement and NK cells. (A) B6 or *C3*^−/−^ mice were treated with an isotype control or an NK1.1 depleting antibody. Four days after antibody administration and confirmation of NK cell depletion, a teratoma assay of HM-KO or HM-KO NK cells was performed. Throughout the course of the experiment, mice were administered antibodies weekly to maintain NK cell depletion. The frequency of detectable teratomas is shown in the top panel, while size of the teratomas is depicted in the bottom panel. The dashed line at 1mm represents the limit of detection. Mean values + SEM are shown for each group of mice. Five mice are represented per group. For teratoma detection, *p<0.05 and ***p<0.001 by log-rank test. For teratoma growth, **p<0.01 and ****p<0.0001 by 2-way ANOVA with post-hoc Tukey’s multiple comparisons test.

## Discussion

Immunological rejection represents the major barrier for organ and bone marrow transplants. This rejection is mediated by multiple arms of the immune system including T cells, NK cells, phagocytes, and complement. Here we described the generation of a line of H1 hESCs ablated for HLA-I and -II, and MICA and B, to prevent T and NK cell-mediated recognition. Additionally, we designed constructs of both mouse and human inhibitory proteins for NK cells, phagocytes, and complement deposition. We demonstrated using teratoma assays that this line evades rejection and persists in immunocompetent mice. These modifications thus prevent xenorejection and may be indicators of barriers that must be overcome in allogeneic hPSC-based therapies. Aside from the obvious requirement that T cell recognition be ablated, we were able to pare down the important remaining modifications to those inhibiting NK cells and complement, while phagocytosis inhibition was not required. We additionally detailed our experience and solutions regarding common pitfalls in genetic modification of hPSCs, including a p53 mutation that was selected for during Cas9 editing, as well as initial silencing of our AAVS constructs, which may serve as an example to others utilizing similar techniques to modify hPSCs. These cells can thus serve as a starting point for stem cell researchers to define transplantation barriers and graft function in immunocompetent animals for therapeutic cell types of interest.

Other recent studies have tested a variety of strategies of editing PSCs to evade aspects of immunological rejection. Combinatorial inhibition of T and NK cells, antigen presenting cells, and macrophages/monocytes in mouse embryonic stem cell (mESC) lines allows them to cross allogeneic barriers^25^. A separate study has shown that removing HLA-A and -B but leaving HLA-C allows human iPSCs (hiPSCs) to evade allogeneic T and NK cells *in vitro* and following adoptive transfer in immunodeficient mice^26^. hPSCs ablated of HLA-I and -II and expressing PD-L1, HLA-G, and CD47 elicit reduced phagocytosis and NK and T cell responses *in vitro* and in CD8^+^ T cell-humanized immunodeficient mice^27^. hESCs engineered to express CTLA4-immunoglobulin and PD-L1 persist in teratoma assays in humanized mice^28^. Deletion of β2m and CIITA and overexpression of CD47 in both mouse and human iPSCs (miPSCs and hiPSCs, respectively) prevents immune rejection upon transplantation in immunocompetent recipients^29^. These same modifications in allogeneic cells were reported to permit cardiomyocyte engraftment to treat heart infarcts in mice^30^, CD19 chimeric antigen receptor T cells to treat CD19^+^ tumors in humanized mice^33^, and pancreatic islets to treat both streptozotocin-induced and autoimmune diabetes in humanized mice^31,32^ or for transplantation into rhesus macaques^31^. In a separate study, PD-L1 overexpression on wild-type hiPSCs was sufficient to allow engraftment of differentiated human islet-like organoids that persisted and controlled streptozotocin-induced diabetes in humanized mice^34^. Together, these studies suggest that there may be multiple paths to evading rejection but also highlight the lack of consensus standards to test immunogenicity of the edited cells and grafts.

No single method to test immunogenicity is perfect, and it is worth considering the key caveats to each type of approach. In our humanized mouse models, despite robust human B and T cell engraftment, even unmodified hESCs were tolerated. While it is possible that alternative humanization systems or recipients would have improved these experiments, none of these models fully recapitulates a normally functioning immune system^117–119^. Thus, the immunological barriers to transplantation may be substantially underestimated in humanized mice. Specific aspects of human responses can be tested *in vitro*, but it is difficult to accurately account for the complex cross talk between immune cells and molecules that would occur in anatomically regulated ways *in vivo*. Another option is to test species-matched allogeneic transplants (i.e. mouse PSCs in mice or nonhuman primate PSCs in nonhuman primates). While this is a strong approach, it does not directly test the modified hPSC products that would ultimately be used therapeutically. We therefore utilized xenotransplants as a readout for immunogenicity with the full acknowledgment of several caveats. For instance, many of the immune evasion proteins in our constructs have mouse and human orthologs, but some do not (i.e. HLA-G and H2-K^b^). However, crossing xenogeneic immune barriers is generally considered to be a higher bar than allogeneic transplantation, even when MHC is mismatched^121–132^. In this xenogeneic system, we were unable to observe a functional effect of CD47, PD-L1, or CD24 overexpression. Perhaps in the background of all the other modifications we made, these changes become redundant, suggesting that there may be multiple ways to achieve the same functional endpoint of a minimally immunogenic PSC line.

A limitation of our study is the exclusive use of teratomas to examine immunogenicity. PSCs and differentiated cells may have different immune barriers based on the intrinsic properties of the cell type and the anatomical and orthotopic site of transplantation^172,173^. Nonetheless, that xenogeneic barriers can be overcome at all in any cell type through genetic modifications lends confidence that development of ‘universal’ hPSC-based therapies is genuinely possible. Moreover, hESC lines, based off well-characterized H1 cells, that can cross mouse xenobarriers can be incorporated rapidly into ongoing translational efforts by other investigators to test in their cells of interest.

## Acknowledgments

This work was supported by the New York Stem Cell Foundation Robertson Investigator Award NYSCF-RI14 (D.B.), NIH R21AI132910 (D.B.), JDRF 3-SRA-2020-895-S-B (P. A-G. and D.B.) and 2-SRA-2017-365-S-B (B.J. and D.B.), and Bill and Melinda Gates Foundation OPP1206188 (D.B.). Additional funding was provided by Lisa Dean Moseley Fellowship (P. A.-G.), the New York Stem Cell Foundation (F.J.M., Jr. and J.W.), and NIH R01CA205239 and P50CA171963 (T.A.F.). We thank the Genome Engineering and iPSC Center at Washington University in St. Louis for generating the HM-KO hESC line.

## Author contributions

Conceptualization and methodology, H.A.P., P. A.-G., B.J., J.P.C., T.A.F., J.P.A., S.M.P.-M., F.J.M., Jr., and D.B.; Formal Analysis, H.A.P., P. A.-G., J.W., J.P.C., S.M.P.-M., F.J.M., Jr., and D.B.; Investigation, H.A.P., P. A.-G., J.W., and J.P.C.; Resources, P. A.-G., T.A.F., J.P.A., S.M.P.-M., F.J.M., Jr., and D.B.; Writing – Original Draft, H.A.P. and D.B.; Writing – Reviewing & Editing, H.A.P., P. A.-G., J.W., J.P.C., T.A.F., J.P.A., S.M.P.-M., F.J.M., Jr., and D.B.; Supervision, S.M.P.-M., F.J.M., Jr., and D.B.; Project Administration, D.B.; Funding Acquisition, D.B., P. A.-G., B.J.

## Declarations of Interests

Sana Biotechnology has licensed intellectual property of H.A.P., D.B., and Washington University in St. Louis. Gilead Biosciences has licensed intellectual property of D.B. and Stanford University. Clade Therapeutics has licensed intellectual property of D.B., H.A.P., and The University of Arizona. D.B. is a co-founder of Clade Therapeutics. D.B. served on an advisory panel for GlaxoSmithKline. T.A.F. has licensed patents, equity, consulting, and potential royalty interests in Wugen; equity interests in Indapta Therapeutics and Orca Bio; and serves as a consultant for Affimed, AI Proteins, Smart Immune, Simcha. B.J. is an employee of AstraZeneca. The authors have no additional financial interests.

## Methods

### hESC Cell Culture

All research was approved by the Embryonic Stem Cell Research Oversight Committees at Washington University in St. Louis and The University of Arizona. H1 hESCs were obtained from WiCell (WA01). hESCs were cultured on Matrigel (Corning) coated plates in mTeSR or mTeSR Plus (STEMCELL Technologies). Cell passaging was performed with Accutase (Innovative Cell Technologies). hESCs were thawed in media containing 10μM Y-27632 (hydrochloride) ROCK inhibitor (Cayman Chemical Company). hESCs were frozen in mFreSR (STEMCELL Technologies). Following fluorescence-activated cell sorting (FACS), hESCs were cultured with CloneR (STEMCELL Technologies) per manufacturer’s instructions. Cultures were maintained at 37°C with 5% CO_2_.

### Generation of HM-KO hESCs

H1 hESCs were pretreated with Revitacell (ThermoFisher) for 1hr, then nucleofected (Lonza Nucleofector 4D, program CA-137) with a Cas9 construct (modified version of pMJ915, a generous gift from Chris Jeans) and up to 3 gRNA-encoding vectors to target each gene. The pool of transfectants was then subjected to next generation MiSeq analysis (Illumina) to estimate the frequency of frameshift mutations in each targeted gene. Individual colonies were picked and screened via Miseq to identify clones with frameshift mutations. Clones with frameshift mutations were then single cell-sorted. Another round of Miseq was performed to verify the mutations and confirm the absence of mosaicism. Clones were then expanded and karyotyped by Cell Line Genetics. The gRNAs and primers used for each targeted gene are listed in **Table S4**. P53 was sequenced with the Accel-Amplicon Comprehensive TP53 Panel (Swift Biosciences) according to the manufacturer’s protocol. To correct the *p53* mutation, HM-KO hESCs were nucleofected with a complex of Cas9 protein bound to a gRNA targeting 5’ GCATGGGCGGCATGAACCAGNGG 3’ along with a correction ssODN 5’ CCTGGAGTCTTCCAGTGTGATGATGGTGAGGATGGGCCTCCTGTTCATGCCGCCCATGCAGGAACTGTTACACATGTAGTTG 3’ and GFP plasmid. Cells were single cell sorted and screened for correction using Miseq with forward primer 5’ AGATCACGCCACTGCACTCCAGCCT 3’ and reverse primer 5’ CGCCGGGGATGTGATGAGAGGTGGA 3’.

### Flow Cytometry

Cells were resuspended in phosphate buffered saline (PBS) with 5% adult bovine serum (ABS) and 2mM ethylenediaminetetraacetic acid (EDTA) prior to staining. hESCs to be sorted were harvested and stained in mTeSR Plus with CloneR. The following human antibodies were purchased from BioLegend: β2-microglobulin (2M2) – APC; TruStain FcX; HLA-A, B, C (W6/32) – APC/Cy7; HLA-E (3D12) – PE; HLA-G (87G) – APC, PE/Dazzle 594; MICA/MICB (6D4) – PE, PerCP/Cy5.5; HLA-DR (L243) – Brilliant Violet 605, Brilliant Violet 650; CD7 (CD7-6B7) – PE; CD34 (581) – PE/Cy7, APC/Cy7; CD4 (A16A1) – PE; CD4 (OKT4) – Brilliant Violet 421, APC; CD8 (SK1) – APC/Cy7; CD14 (M5E2) – PE/Cy7; CD13 (WM15) – PE/Cy7; CD43 (CD43-10G7) – APC, PE/Cy7; CD45RA (HI100) – Alexa Fluor 700; CD45 (2D1) – APC; CD73 (AD2) – PE; CD1c (L161) – Brilliant Violet 510; CD3 (OKT3) – PerCP/Cy5.5; CD10 (HI10a) – PerCP/Cy5.5; CD11b (ICRF44) – Brilliant Violet 650; CD19 (HIB19) – Brilliant Violet 421; CD19 (SJ25C1) – Brilliant Violet 510; CD33 (WM53) – PerCP/Cy5.5; CD34 (561) – FITC; CD38 (HIT2) – PE/Cy7, biotin; CD46 (TRA-2-10) – APC/Cy7, PE; CD47 (CC2C6) – PE/Cy7; CD49f (GoH3) – Brilliant Violet 421; CD49d (9F10) – PE/Cy5; CD55 (JS11) – APC; CD56 (5.1H11) – APC; CD56 (HCD56) – PE/Cy7; CD59 (H19) – FITC; CD85j (GHI/75) – biotin; CD90 (5E10) – APC; CD107a (H4A3) – Brilliant Violet 421; CD141 (M80) – PE; CD158d (mAb 33 (33)) – PE; CD184 (12G5) – Brilliant Violet 421; CD135 (BV10A4H2) – biotin; TCR γ/d (B1) -PE; CD11c (Bu15) – Alexa Fluor 700. The following human antibodies were purchased from BD: CD11b (ICRF44) – Alexa Fluor 488; CD16 (3G8) – APC-Cy7; CD33 (HIM3-4) – FITC; CD46 (E4.3) – FITC; CD59 (p282 (H19)) – PE. Human HLA-ABC (W6/32) – PE was purchased from Invitrogen. Human CD159a (REA110) – FITC was purchased from Miltenyi Biotec. The following mouse antibodies were purchased from BioLegend: CD3 (145-2C11) – Brilliant Violet 510, PerCP/Cy5.5; CD4 (RM4-5) – Brilliant Violet 605; CD8 (53-6.7) – Alexa Fluor 700; CD19 (6D5) – Brilliant Violet 421; CD45 (30-F11) – Alexa Fluor 488; B220 (RA3-6B2) – FITC; NK-1.1 (S17016D) – PE; Ly-6G/Ly-6C (RB6-8C5) – APC; CD47 (miap301) – FITC, PE, Brilliant Violet 421, APC/Cy7, PE/Dazzle 594; CD55 (RIKO-3) – PE, PE/Cy7; CD59 (mCD59.3) – PE; CD274 (10F.9G2) – PE, Brilliant Violet 605, Brilliant Violet 650; H-2Kb (AF6-88.5) – Alexa Fluor 647, FITC, Brilliant Violet 421; CD29 (HMβ1-1) – APC/Cyanine7, Alexa Fluor 488; Ig light chain k (RMK-45) – FITC; CD21/35 (7E9) – APC; CD24 (M1/69) – Alexa Fluor 700, Brilliant Violet 510, Pacific Blue. The following mouse antibodies were purchased from BD: CD11b (M1/70) – PE/Cy7; Crry/p65 (1F2) – biotin, BV786; CD24 (M1/69) – PE/Cy7. The following mouse antibodies were purchased from Miltenyi Biotec: CD55 – biotin; Qa-1b (6A8.6F10.1A6) – APC; Qa-1b – biotin. Mouse IgG – UNLB was purchased from Southern Biotech. Streptavidin – BV605, BV421, PE-Cy7 were purchased from BD. DAPI and propidium iodide were purchased from Sigma-Aldrich. Cells were analyzed on a BD LSR II, BD Fortessa, BD Fortessa X-20, or Cytek Aurora. All fluorescence activated cell sorting was performed on a BD FACS Aria II or III. Data was analyzed using FlowJo software (FlowJo Enterprise). hESCs were treated with PBS as a vehicle control or 20ng/mL hIFNγ (PeproTech) for 24hrs to induce HLA-I expression.

### DC-Like Cell Differentiations

hESCs were differentiated into hematopoietic progenitors using either an embryoid body culture described previously^174^ or the STEMdiff Hematopoietic Kit (STEMCELL Technologies) per the manufacturer’s instructions. Hematopoietic progenitors were collected then cultured in flasks coated with 20mg/mL poly-HEMA (Sigma-Aldrich) in α-MEM with 10% fetal bovine serum (FBS), Glutamax, penicillin/streptomycin, and 100ng/mL hGM-CSF (PeproTech) for 8-10d with half medium changes every 4d^175^. Cells were collected and spun over 20% Percoll (GE Healthcare). The cells at the interface were collected and cultured in poly-HEMA-coated flasks with StemSpan SFEM (STEMCELL Technologies) supplemented with lipid mixture 1 (Sigma-Aldrich), 100ng/mL hGM-CSF (Peprotech), and 100ng/mL hIL-4 (PeproTech) for 7-9d with half medium changes every 4d. Cells were then collected and cultured in poly-HEMA-coated flasks in StemSpan SFEM with lipid mixture 1 and 400ng/mL A23187 calcium ionophore (Sigma-Aldrich) for 2d. Human PBMCs were used as a control following culture in RPMI supplemented with 10% FBS, 2-mercaptoethanol, penicillin/streptomycin, 100ng/mL hGM-CSF, and 100ng/mL hIL-4 for 7-10d.

### Whole Exome Sequencing

DNA was isolated using a Genomic DNA Mini Kit (IBI Scientific). Whole exome sequencing was performed by the University of Chicago Genomics Facility. Sequencing data was first processed using the BWA pipeline from the Broad Institute^176–179^ and then by the Ensembl Variant Effect Predictor^180^. Concerning mutations were identified using the Human Clinical Variation Database^181^. Raw sequencing data is available at the NCBI Sequence Read Archive (accession PRJNA987027).

### Mouse Lines

All animal procedures used in this study were approved by the Animal Care and Use Committees at Washington University in St. Louis and the University of Arizona. NOD.Cg-*Kit^W-41J^ Tyr* ^+^ *Prkdc^scid^ Il2rg^tm1Wjl^*/ThomJ (NSG-W41) mice were obtained from The Jackson Laboratory (stock number 026622)^113–115^. C57BL6/N mice were obtained from Charles River Laboratories. B6.129S4-*C3^tm1Crr^*/J (*C3^−/−^*) mice were obtained from The Jackson Laboratory (stock number 029661)^171^.

### Humanized Mice

Human cord blood was obtained from the St. Louis Cord Blood Bank. CD34^+^ cells were enriched using Dynabeads CD34 Positive Isolation Kit (Life Technologies) per the manufacturer’s instructions. 1 − 10^5^ cells were injected retro-orbitally into unconditioned NSG-W41 recipients. At least 2mo post-transplantation, mice were bled to confirm human chimerism. Peripheral blood was collected in 10mM EDTA/PBS via tail venipuncture of warmed mice. Red blood cells were lysed with 0.15M NH_4_Cl, 10mM KHCO_3_, 0.1mM EDTA, pH 7.2 solution (ACK) prior to antibody staining. For splenic chimerism, spleens were harvested and dissociated using frosted glass microscope slides. Red blood cells were lysed using ACK. Cells were filtered through 70μm nylon mesh prior to antibody staining.

### Teratoma Assays

hESCs were collected from culture and resuspended at 10 − 10^6^ cells/mL in a 1:1 mix of Matrigel and PBS. 100µL of cells were injected subcutaneously into the hind flank of each mouse. Teratoma growth was monitored weekly, and diameters were measured with a caliper once the diameter was confidently measurable (∼1.2mm).

### Lentivirus Construct Design, Production, and Transduction

Immune evasion genes were cloned without stop codons into lentiviral vectors downstream of a ubiquitin promoter and in-frame with a downstream T2A-GFP cassette (base vector was a gift from A. Bredemeyer, Washington University)^182^. Lenti-X 293T cells (Takara Bio USA) were cultured at 37°C with 5% CO_2_ in DMEM with 10% FBS, nonessential amino acids, Glutamax, sodium pyruvate, and penicillin/streptomycin. Cells were transfected at ∼60% confluency in 10cm^2^ tissue culture plates using 30μL GeneJuice Transfection Reagent (Sigma-Aldrich) with 5μg of lentiviral vector, 3.25μg psPax2 (Addgene 12260), and 1.75μg VSV.G (Addgene 12259). Medium was changed 6-8hr post-transfection, and viral supernatant was harvested 48 and 72hr later. 12mL of viral supernatant was mixed with 3mL of 25% polyethylene glycol 8000 (Sigma-Aldrich) in PBS and incubated overnight at 4°C. This mixture was centrifuged at 3,000 x g for 20 min, supernatant was discarded, and the pellet was resuspended in 100μL PBS. Aliquoted lentivirus was stored at −80°C until time of use. 5-10μL was used to transduce one well of hESCs in a six well plate. Transduced cells were sorted for GFP and protein expression.

### hAAVS Construct Design and hESC Transfection

AAVS constructs were gene-synthesized by BioBasic. AAVS constructs were transfected into hESCs along with LentiCRISPRv2-mCherry (Addgene plasmid 99154), encoding a gRNA for human AAVS1 targeting (GGAAGAGAGTAGGTCGAAG) using GeneJuice Transfection Reagent (Sigma-Aldrich). Transfected cells were selected with either 0.5μg/mL puromycin or 100μg/mL neomycin. Transfected populations were also purified using FACS.

### Immunohistochemistry (IHC)

Teratomas were harvested and fixed in 4% paraformaldehyde solution (Santa Cruz Biotechnology, Inc.) for 24hr at room temperature then washed with PBS 3 times and stored in PBS at 4°C until ready for use. Samples were cryoprotected with sucrose (Sigma-Aldrich), cryopreserved in Tissue-Tek O.C.T. compound (Sakura), and sectioned at 5µm with a Leica CM3050 S cryostat. Sections were incubated with PBS containing 0.1% triton X-100 (Sigma-Aldrich) and 5% donkey serum (Jackson ImmunoResearch). Sections were incubated overnight with unconjugated goat anti-mouse/rat CD47 N-terminal IgV-like extracellular domain antibody (R&D Systems), biotin-conjugated anti-mouse CD55 REAfinity antibody (clone REA300, Miltenyi Biotec), unconjugated rabbit anti-mouse CD59a antibody (clone 108, SinoBiological), and AF594-conjugated anti-human mitochondria antibody (clone 113-1, Novus Biologicals) followed by a 1hr incubation with Alexa Fluor 647-conjugated AffiniPure donkey anti-goat IgG antibody (Jackson ImmunoResearch), Alexa Fluor 647-conjugated streptavidin (Jackson ImmunoResearch), or Alexa Fluor Plus 555-conjugated donkey anti-rabbit IgG antibody (Invitrogen). Slides were mounted with ProLong Gold Antifade Mountant with DNA stain DAPI (Invitrogen) and imaged on a Zeiss LSM 800 confocal laser scanning microscope. The resulting images were processed and analyzed with Fiji (ImageJ).

### Antibody Depletion

Mice were bled (as described in the humanized mice section) prior to the first injection of depleting antibodies. Mice were injected intraperitoneally with 250µg of one of the following depleting antibodies: mouse IgG2a (clone C1.18.4; BioXCell), rat IgG2b (BioXCell), CD8 (clone YTS169.4; BioXCell), CD4 (clone GK1.5; BioXCell), CD20 (clone SA271G2; BioLegend), or NK1.1 (clone PK136; BioXCell). Four days after injection, mice were bled to confirm depletion. hESCs were transplanted into these mice 5d after the initial depleting antibody injection. Depleting antibodies were administered weekly throughout the teratoma assay.

### Chinese Hamster Ovary Cell Culture

Chinese hamster ovary (CHO) cells were cultured in DMEM with 10% FBS and penicillin/streptomycin. Cells were maintained at 37°C with 5% CO_2_. Human AAVS constructs were transfected into CHO cells using the Gene Pulser MXcell Electroporation System (Bio-Rad). Transfected cells were selected for with 10µg/mL puromycin or 1mg/mL neomycin. Cells expressing the immune evasion proteins were also purified via FACS.

### Complement Deposition Assay

CHO cells were incubated with 1µg/mL anti-CHO antibody (Cygnus Technologies) for 30min at 4°C. Cells were washed with either GVB++ or GVB° + MgEGTA (Complement Technology) for the classical or alternative pathway, respectively. Cells were resuspended in 10% C7-deficient serum (Sigma-Aldrich) in the appropriate GVB buffer and incubated at 37°C for 45min with shaking. Cells were washed with 1% FBS/PBS then stained with mouse anti-human C3c, C3d, C4c, C4d, or C5 antibody (Quidel) for 30min at 4°C. Cells were washed with 1% FBS/PBS then stained with PE/Cy7-conjugated rat anti-mouse Igκ light chain antibody (BD) for 30min at 4°C. Cells were washed and blocked with unlabeled mouse IgG (Southern Biotech) prior to subsequent staining and analysis.

### K562 Cell Culture

K562s were cultured in DMEM with 10% FBS, Glutamax, nonessential amino acids, sodium pyruvate, and penicillin/streptomycin. Cultures were maintained at 37°C with 5% CO_2_. Human AAVS constructs were transfected into K562s using the Gene Pulser MXcell Electroporation System (Bio-Rad). Transfected cells were selected for with 10µg/mL puromycin or 1mg/mL neomycin. Cells expressing the immune evasion proteins were also purified via FACS.

### NK Cell CD107a Degranulation Assay

PBMCs were plated in RPMI with 10% FBS, Glutamax, nonessential amino acids, sodium pyruvate, HEPES, and penicillin/streptomycin supplemented with 1ng/mL hIL-15 (PeproTech) and cultured overnight at 37°C with 5% CO_2_. PBMCs were either cultured alone (unstimulated) or at a 10:1 ratio with K562s. Cells were stained with CD107a for 1hr at 37°C. GolgiStop/Plug (BD) was then added per manufacturer’s instructions and cells were incubated for 5hr at 37°C. Cells were washed and stained for analysis.

## Statistics

All statistical analyses were conducted using GraphPad Prism and are detailed in each figure legend.

## Figures

Figures were created with BioRender.com.

## Supplemental Figure Legends

**Figure S1, related to Figure 1.**
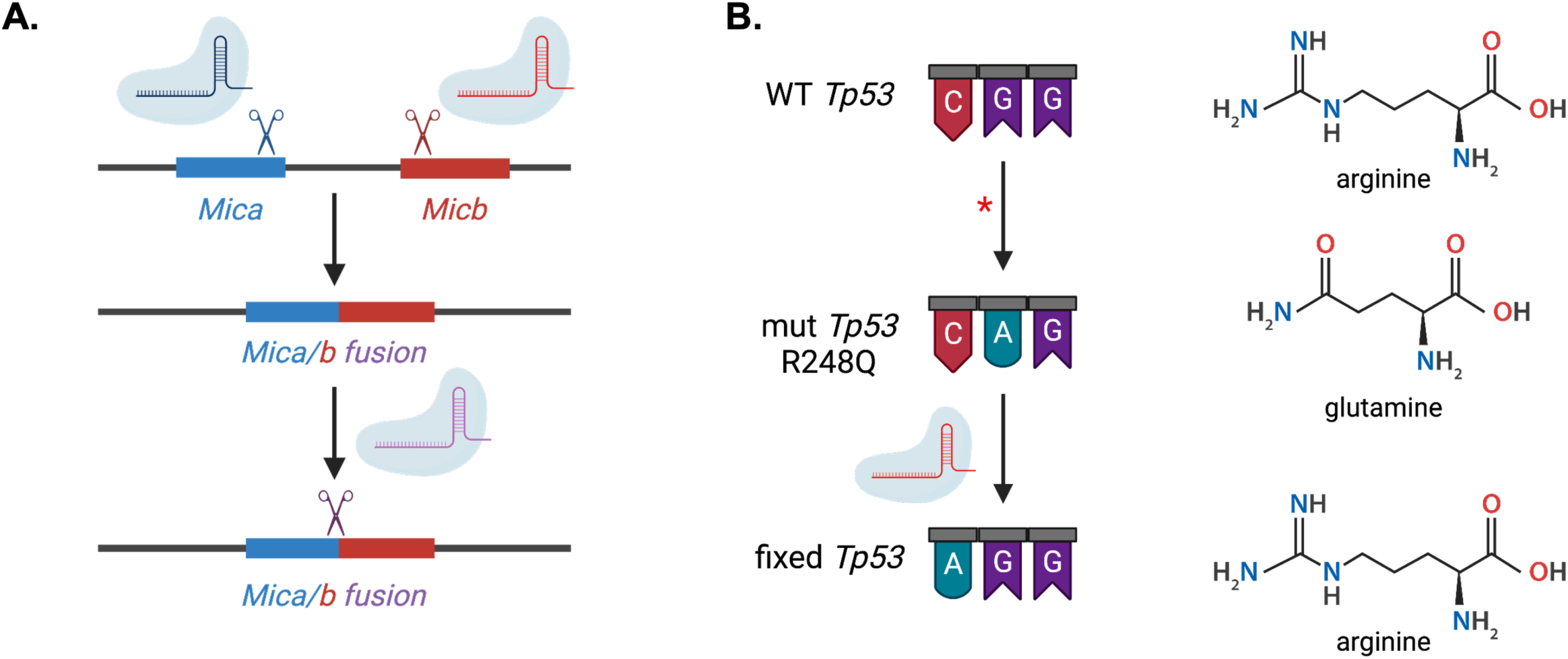
Ablation of the MICA/MICB locus and correction of a p53 mutation in HLA-deficient hESCs. (A) Schematic of the *Mica/b* fusion generated in one allele. gRNAs for *Mica* and *Micb* cut their respective genes but left behind an in-frame fusion of *Mica* and *Micb* in one allele. This fusion was edited out of frame with a different gRNA. (B) Schematic of the R248 residue of wild-type *Tp53*, the identified mutation, and the reverted line. Wild-type p53 encodes an arginine (CGG) at position 248. In the first step of generating the HM-KO line, clones were inadvertently selected that carried a mutated CAG encoding glutamine. Using a gRNA and Cas9, this codon was reverted to an arginine-encoding AGG sequence.

**Figure S2, related to Figure 3.**
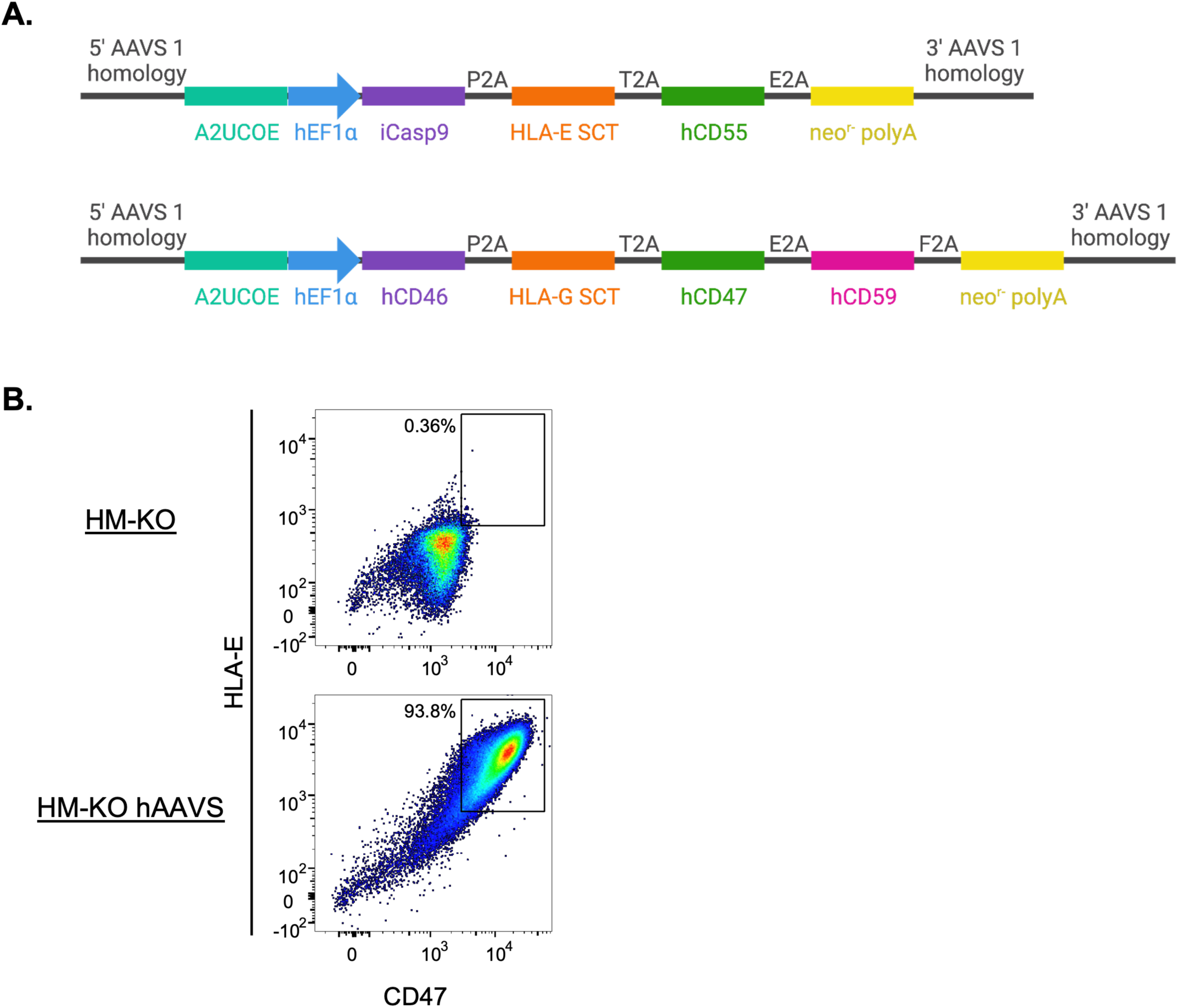
Human immune evasion construct design and expression. (A) Two human AAVS constructs contain the inhibitory genes of interest. Each construct contains an A2UCOE insulator element, and expression is driven by the human elongation factor 1α (hEF1α) promoter. The immune evasion genes HLA-E SCT, hCD55, hCD46, HLA-G SCT, hCD47, and hCD59 are linked by 2A sequences. A suicide gene (iCasp9) is present in one construct, and both contain a neomycin resistance gene. The constructs have 5’ and 3’ AAVS1-targeting homology arms. (B) Flow cytometric analysis of human inhibitory protein expression on HM-KO cells, as well as HM-KO cells transfected with both human AAVS constructs (HM-KO hAAVS).

**Figure S3, related to Figure 3.**
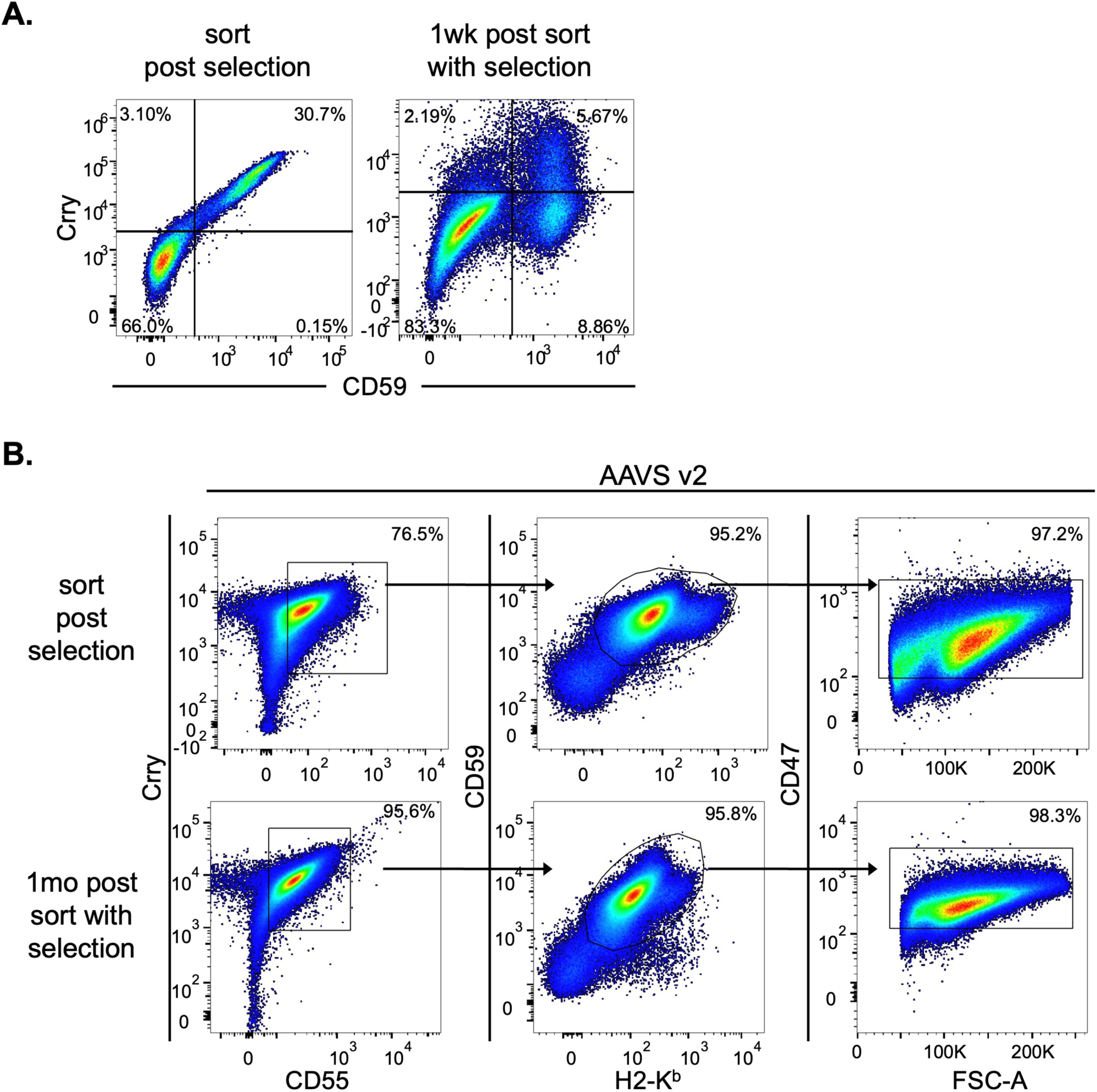
Modifying AAVS targeting site reduces silencing of expression. (A) Sample flow cytometry plots of immune evasion expression over time in HM-KO hESCs transfected with the original mouse AAVS constructs. Expression is shown following drug selection of transfected cells prior to sorting of Crry^+^ CD59^+^ double positive cells. Expression was then analyzed a week after sorting and further drug selection. (B) Sample flow cytometry plots of immune evasion expression over time in HM-KO hESCS transfected with the redesigned AAVS constructs. The top row shows consecutive gating of transfected cells following drug selection. These cells were sorted and kept on selection. Two months later expression was analyzed again as shown in the bottom row.

**Figure S4, related to Figure 4.**
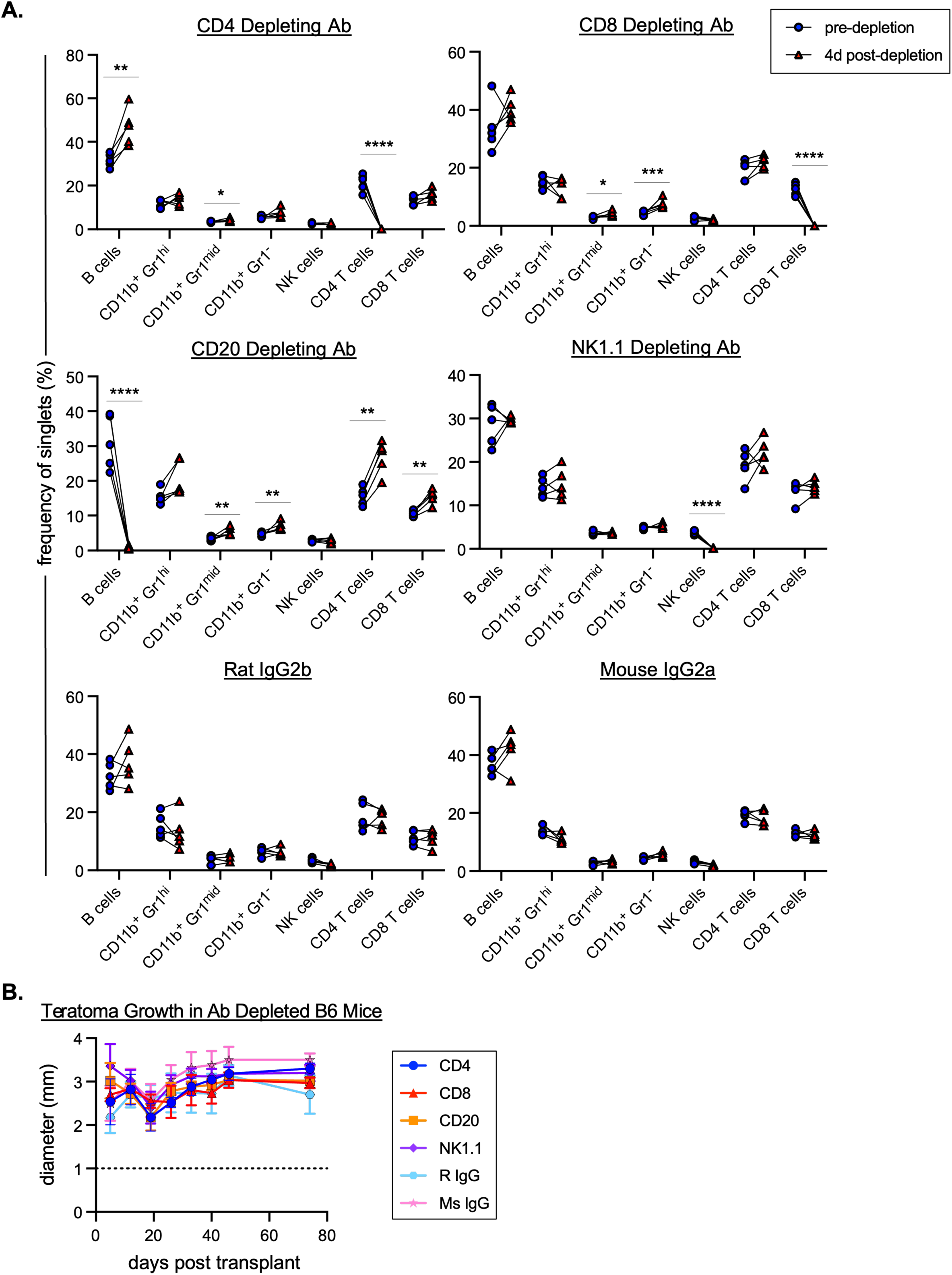
Depletion of T cells, B cells, and NK cells does not enhance HM-KO p53_mut_ lenti teratoma growth. (A) Confirmation of depletion of immune cells following antibody treatment. B6 mice were injected with CD4, CD8, CD20, or NK1.1 depleting antibodies or with control rat IgG2b or mouse IgG2a. Prior to injection of antibodies, mice were bled to determine the baseline levels of circulating immune cells. Four days following antibody injection, mice were bled to quantify the extent of depletion. Five wild-type mice are in each depleting antibody group, and pre-depletion and post-depletion frequencies are paired for each individual mouse. *p<0.05, **p<0.01, ***p<0.001, and ****p<0.0001 by Student’s 2-tailed paired t-test. (B) Teratoma assay to assess growth of HM-KO p53_mut_ lenti hESCs in B6 mice treated with depleting antibodies. Mice were injected with depleting antibodies 5d prior to the injection of HM-KO p53_mut_ lenti hESCs. Throughout the assay, mice were injected weekly to maintain the depletion of each cell type. Mean values + SEM are shown for each group of mice. Five mice are represented per group. P values greater than 0.05 by 2-way ANOVA with post-hoc Tukey’s multiple comparisons test are not depicted.

**Figure S5, related to Figure 4.**
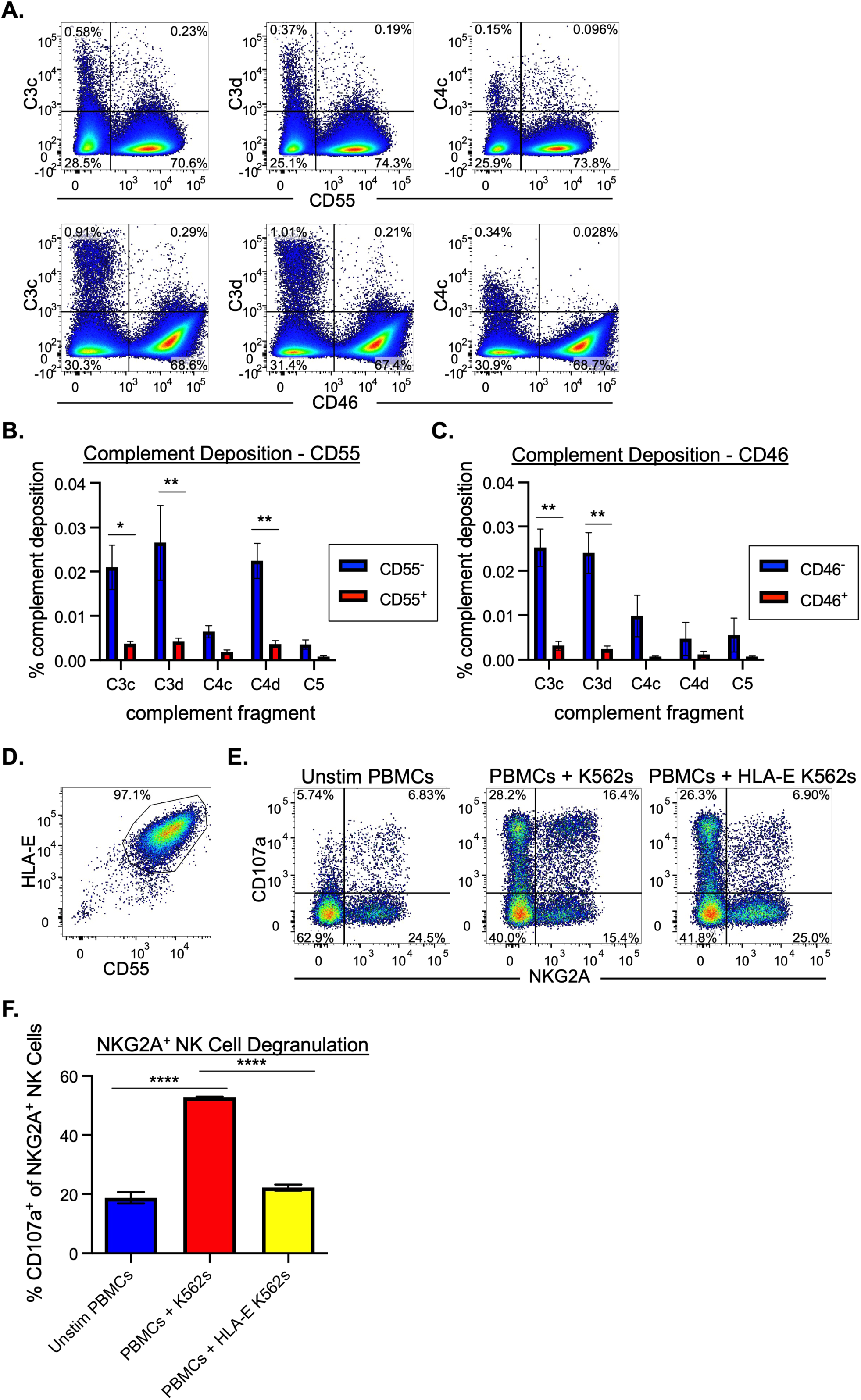
Human CD55 and CD46 inhibit complement deposition, while HLA-E inhibits NKG2A^+^ NK cell degranulation. (A) Representative flow cytometry plots of complement deposition on Chinese hamster ovary (CHO) cells with or without the expression of CD55 or CD46. CHO cells were transfected with the human AAVS constructs (Figure S2) containing either human CD55 (top row) or CD46 (bottom row). Cells were incubated with anti-CHO antibody followed by human C7-deficient serum and were stained for complement fragment deposition (C3c, C3d, C4c). (B-C) Quantification of complement deposition on cells expressing CD55 (B) or CD46 (C). Mean values ± SEM are shown for 3-7 replicates from two independent experiments. *p<0.05 and **p<0.01 by Student’s 2-tailed t-test. (D) Flow cytometry plot of K562s transfected with the human AAVS construct containing HLA-E and hCD55. Cells were transfected, placed under selection, and sorted to obtain a pure population for downstream NK cell degranulation assays. (E) Representative flow cytometry plots of NK cell degranulation assays utilizing unstimulated PBMCs, PBMCs mixed with K562s, and PBMCs mixed with HLA-E-expressing K562s. Degranulation is measured by the expression of CD107a. Populations shown have been gated on CD56^+^ NK cells. (F) NKG2A^+^ NK cell degranulation assay. The frequency of CD107a-expressing cells of NKG2A^+^ NK cells was analyzed from unstimulated PBMCs, PBMCs with K562s, and PBMCs with HLA-E^+^ K562s. Mean values ± SEM are shown for 6 replicates from two independent experiments. ****p<0.0001 by Student’s 2-tailed t-test.

**Figure S6, related to Figure 5.**
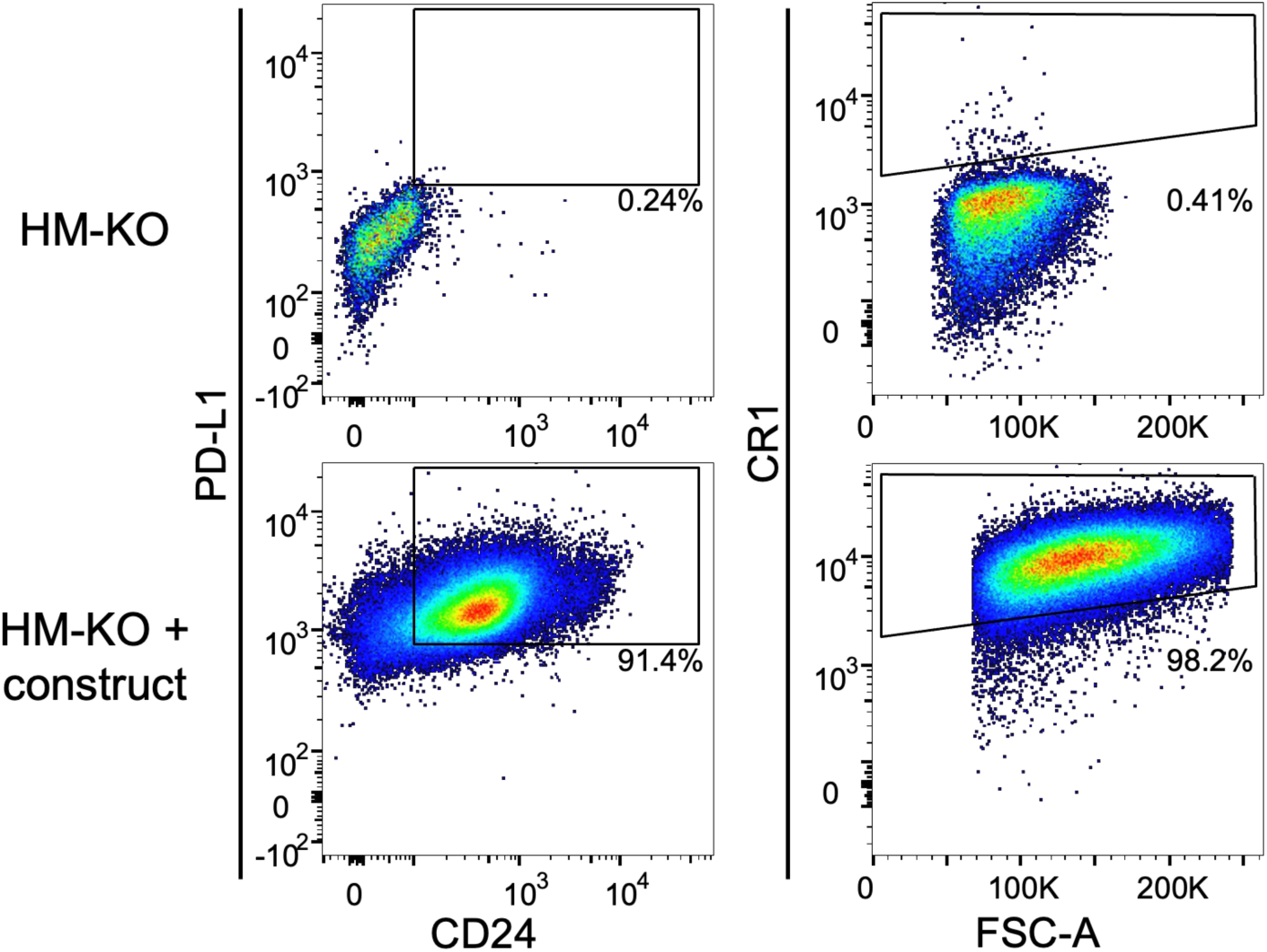
Expression of PD-L1, CD24, and CR1. Flow cytometric analyses of the expression of PD-L1, CD24, and CR1 on HM-KO cells and HM-KO cells transduced with PD-L1 and CD24 lentiviruses or transfected with a CR1 AAVS construct.

## Supplemental Tables

**Table S1.** Indel data for the 6 genes ablated to generate the HM-KO hESC line. A list of the insertions and deletions (indels) found for each gene ablated and the read count frequency of each edited allele.

| Targeted gene | Indel | Reads | Total reads (%) |
| --- | --- | --- | --- |
| *β2m* | -1 | 5032 | 52.9 |
|  | 5 | 4454 | 46.9 |
|  | -2 | 11 | 0.1 |
|  | 4 | 5 | 0.1 |
| *Tap1* | -2 | 1300 | 52.5 |
|  | 1 | 1172 | 47.3 |
|  | -3 | 2 | 0.1 |
|  | 0 | 1 | 0.0 |
| *Cd74* | -10 | 787 | 48.9 |
|  | -8 | 756 | 47.0 |
|  | 0 | 31 | 1.9 |
|  | -41 | 17 | 1.1 |
| *Ciita* | -2 | 456 | 51.4 |
|  | 2 | 422 | 47.6 |
|  | 0 | 9 | 0.9 |
|  | 1 | 1 | 0.1 |
| *Mica* | -1 | 1500 | 97.3 |
|  | -2 | 39 | 2.5 |
| *Micb* | 1 | 1064 | 98.0 |
|  | 0 | 22 | 2.0 |
| *Mica/b fusion* | -13 | 460 | 95.8 |

**Table S2.** hESC lines generated. Modifications and nomenclature of edited hESC lines.

| Name | Knocked out genes | Added genes (method) | Other notes |
| --- | --- | --- | --- |
| H1 | -- | -- | wild-type |
| HLA-I/II-KO | *β2m*, *Tap1*, *Cd74*, *Ciita* | -- | *Tp53* mutation |
| HM-KO p53<sub>mut</sub> | *β2m*, *Tap1*, *Cd74*, *Ciita*, *Mica*, *Micb* | -- | *Tp53* mutation |
| HM-KO | *β2m*, *Tap1*, *Cd74*, *Ciita*, *Mica*, *Micb* | -- | *Tp53* mutation fixed |
| HM-KO p53<sub>mut</sub> lenti | *β2m*, *Tap1*, *Cd74*, *Ciita*, *Mica*, *Micb* | *Crry*, *mCd55*, *mCd59*, *H2-K<sup>b</sup>*, *mCd47* (lentivirus) | *Tp53* mutation fixed |
| HM-KO lenti | *β2m*, *Tap1*, *Cd74*, *Ciita*, *Mica*, *Micb* | *Crry*, *mCd55*, *mCd59*, *H2-K<sup>b</sup>*, *mCd47* (lentivirus) | *Tp53* mutation fixed |
| HM-KO AAVS | *β2m*, *Tap1*, *Cd74*, *Ciita*, *Mica*, *Micb* | *Crry*, *mCd55*, *mCd59*, *H2-K<sup>b</sup>*, *mCd47*, *Qa1* (AAVS construct) | *Tp53* mutation fixed |
| HM-KO comp | *β2m*, *Tap1*, *Cd74*, *Ciita*, *Mica*, *Micb* | *Crry*, *mCd55*, *mCd59* (lentivirus) | *Tp53* mutation fixed |
| HM-KO NK | *β2m*, *Tap1*, *Cd74*, *Ciita*, *Mica*, *Micb* | *H2-K<sup>b</sup>*, *Qa1* (lentivirus) | *Tp53* mutation fixed |
| HM-KO phago | *β2m*, *Tap1*, *Cd74*, *Ciita*, *Mica*, *Micb* | *mCd47*, *mCD24*, *mPdl1* (lentivirus) | *Tp53* mutation fixed |
| HM-KO CR1 | *β2m*, *Tap1*, *Cd74*, *Ciita*, *Mica*, *Micb* | *mCr1* (AAVS construct) | *Tp53* mutation fixed |
| HM-KO CR1 comp | *β2m*, *Tap1*, *Cd74*, *Ciita*, *Mica*, *Micb* | *mCr1* (AAVS construct), *mCd59* (lentivirus) | *Tp53* mutation fixed |
| HM-KO NK and comp | *β2m*, *Tap1*, *Cd74*, *Ciita*, *Mica*, *Micb* | *Crry*, *mCd55*, *mCd59*, *H2-K<sup>b</sup>*, *Qa1* (lentivirus) | *Tp53* mutation fixed |
| HM-KO CR1 NK and comp | *β2m*, *Tap1*, *Cd74*, *Ciita*, *Mica*, *Micb* | *mCr1* (AAVS construct), *mCd59*, *H2-K<sup>b</sup>*, *Qa1* (lentivirus) | *Tp53* mutation fixed |

**Table S3.** Immune evasion proteins. Human and mouse inhibitory factors in immune evasion constructs and the cell type or pathway that they inhibit.

| Human proteins | Mouse proteins | Inhibits |
| --- | --- | --- |
| CD46 (MCP) | Crry | Complement/C3b and C4b |
| CD55 (DAF) |  | Complement/C3 and C5 convertases |
| CD59 |  | Complement/membrane attack complex |
| HLA-E | Qa1 | NKG2A<sup>+</sup> NK cells |
| HLA-G | - | ILT2/KIR2DL4<sup>+</sup> NK cells |
| - | H2-K<sup>b</sup> | Ly49C<sup>+</sup> NK cells |
| CD47 | | Phagocytes (SIRPα) |
| CD24 |  | Phagocytes (Siglec-10) |
| PD-L1 |  | Phagocytes (PD-1), antigen presentation, T cell activation |
| CR1 |  | Complement/C3b and C4b, C3 and C5 convertases |

**Table S4.**
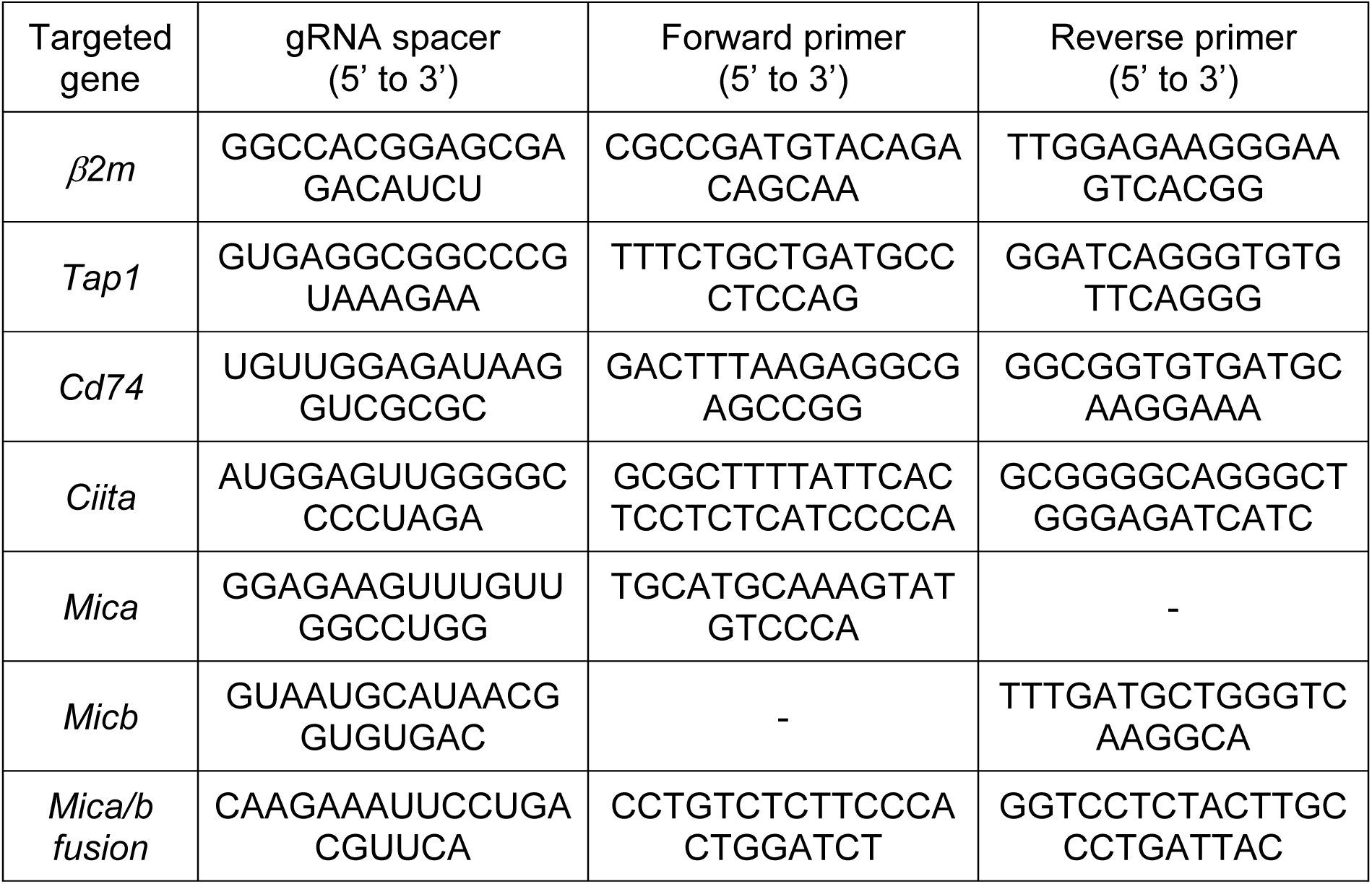
Gene targeting to generate the HM-KO hESC line. gRNA spacers and primers used for gene targeting of H1 hESCs to generate HM-KO hESCs.

## Notes

https://www.ncbi.nlm.nih.gov/bioproject/PRJNA987027/

